# Diseased human pancreas and liver microphysiological system for preclinical diabetes research

**DOI:** 10.1101/2023.07.03.547412

**Authors:** Sophie Rigal, Belén Casas, Kajsa P. Kanebratt, Charlotte Wennberg Huldt, Lisa U. Magnusson, Erik Müllers, Fredrik Karlsson, Maryam Clausen, Sara F. Hansson, Rasmus Jansson Löfmark, Carina Ämmälä, Uwe Marx, Peter Gennemark, Gunnar Cedersund, Tommy B. Andersson, Liisa K. Vilén

## Abstract

Current research on metabolic disorders such as type 2 diabetes relies on animal models because multi-organ diseases cannot be well studied with the standard *in vitro* assays. Here, we connect models of key metabolism organs, pancreas and liver, on a microfluidic chip to enable diabetes research in a human-based preclinical system. Aided by mechanistic mathematical modelling, we developed a two-organ microphysiological system (MPS) that replicates clinically-relevant phenotypes of diabetic dysregulation both in the liver and pancreas compartments. Exposure to hyperglycemia and high cortisone created a diseased pancreas-liver MPS which displayed beta-cell dysfunction, steatosis, elevated ketone-body secretion, increased glycogen storage, and upregulated gluconeogenic machinery. In turn, normoglycemia and physiological cortisone concentration maintained glucose tolerance and stable liver and beta-cell functions. This method was evaluated for repeatability in two laboratories and was effective in multiple pancreatic islet donors. The model also provides a platform to identify new therapeutic targets as demonstrated with a liver-secreted IL-1R2 protein that induced islet proliferation.

## INTRODUCTION

The growing epidemic of type 2 diabetes (T2D) is one of the major medical challenges today. T2D is characterized by hyperglycemia which is caused by dysfunctional communication between several glucose-regulating organs. Understanding the mechanisms of glucose dysregulation is essential for discovering and evaluating effective treatments to prevent or to cure T2D. In healthy individuals, pancreatic beta cells respond to increased blood glucose concentration by secreting insulin. Within minutes, insulin induces the uptake and storage of glucose in the liver and other target organs to restore the normoglycemic glucose concentration in blood^1^ (**Fig. 1A**). Especially, the liver has a central role in glucose homeostasis because it stores glucose in form of glycogen or lipids (*de novo* lipogenesis) during hyperglycemia and produces glucose during hypoglycemia by gluconeogenesis to normalize the glucose concentration^2^. Glucose dysregulation occurs when the target organs become increasingly resistant to insulin and fail to control the blood glucose concentration properly (**Fig. 1B**). Insulin resistance, in turn, evokes increased insulin secretion to compensate for the impaired insulin sensitivity (beta-cell adaptation) and may ultimately result in pancreatic beta-cell failure and overt T2D^3^ (**Fig. 1C**). As T2D is a multi-organ disease, preclinical studies of disease progression mechanisms are currently only possible in animal models. However, animal models used in diabetes research are genetically and physiologically different from humans leading to inaccurate translation^4^. Animal models are, for example, not suitable for studying human-specific new therapeutic modalities^5^ because such drugs directly inhibit disease-causing genes and can have low cross-reaction to the corresponding genes in animals^6^.

**Fig. 1.**
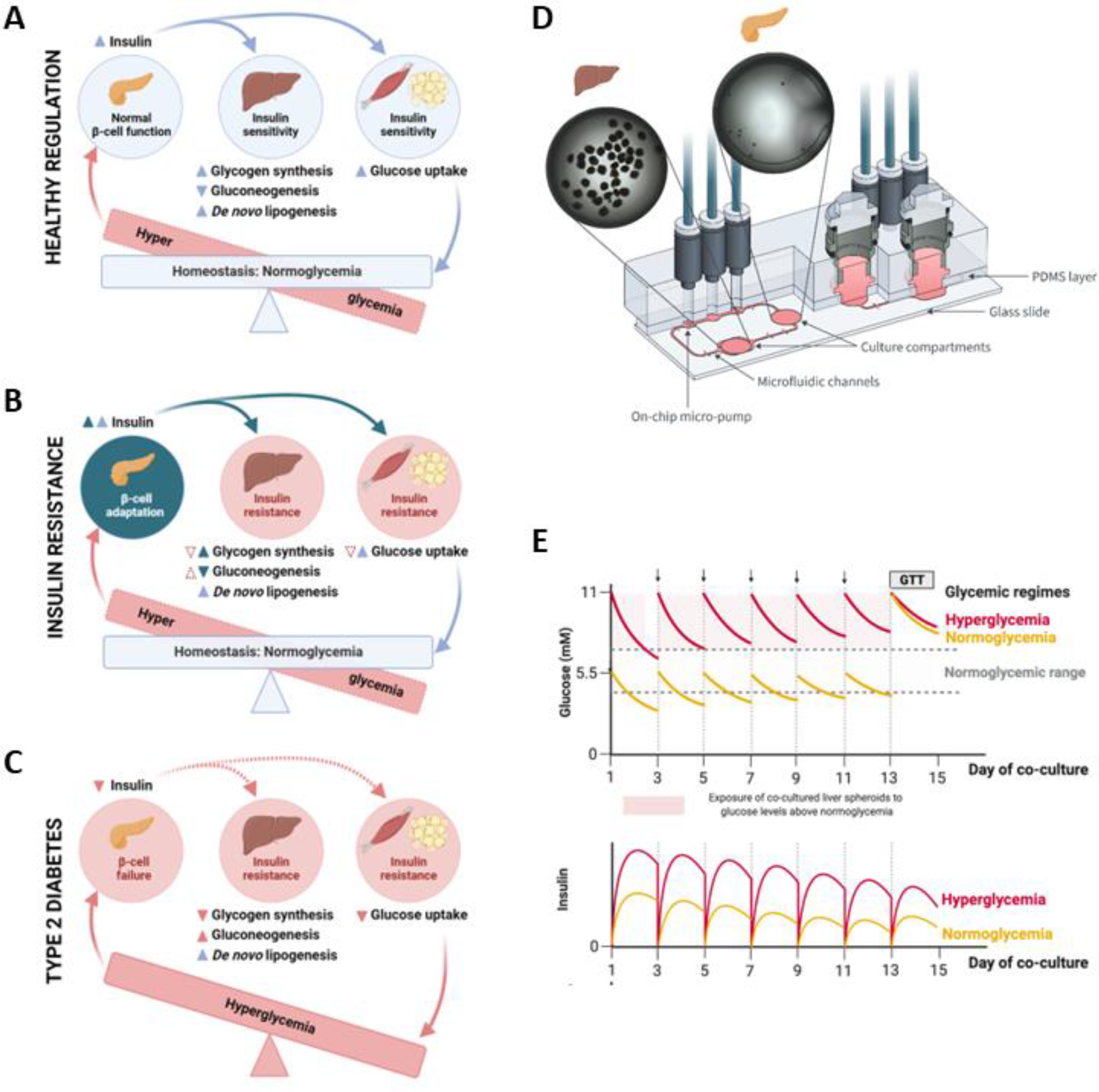
Pancreas-liver MPS for investigation of diabetic glucose dysregulation. (**A**) Healthy glucose regulation. Pancreatic islets prevent long-term hyperglycemia by secreting insulin which promotes glucose uptake and storage as well as de novo lipogenesis 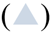 and inhibits glucose release 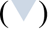 from insulin-sensitive tissues including the liver, muscle, and adipose tissue. (**B**) Glucose regulation in the insulin-resistant state. Insulin resistance causes decreased glucose uptake and storage 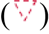 as well as increased glucose release 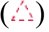 while insulin-stimulated lipogenesis remains unaffected 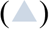. Adaptive insulin secretion 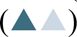 prevents hyperglycemia by normalizing glucose uptake and storage 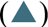 and the inhibition of glucose release 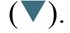. (**C**) Glucose regulation in type 2 diabetes (T2D). Long-term hyperglycemia develops due to a reduced insulin secretion 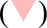 which reduces glucose uptake and storage 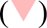 and increases glucose release 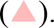. (**D**) Schematic of the pancreas-liver co-culture in the Chip2 microphysiological system. Brightfield images show 40 HepaRG/HHSteC liver spheroids in the outer culture compartment and 10 islets in the inner culture compartment of one Chip2 circuit on day 1 of co-culture. (**E**) Graphical illustrations of representative, previously reported, glucose and insulin responses in the pancreas-liver co-culture^11^ visualize the development of insulin-resistance over time. Arrows indicate medium exchange. GTT; Glucose tolerance test.

The recent advancements in microphysiological systems (MPS) or organ-on-chip models have enabled human *in vitro* studies of physiological organ crosstalk, disease development, and pharmacological effects^7, 8^. Since the pancreas and the liver are central organs in blood glucose regulation, we and others have shown that functional coupling of pancreatic and liver organ models on chip can recapitulate human-relevant pancreas-liver axis^9–11^. In these two-organ models, human islet microtissues (InSphero)^9, 11^ (**Fig. 1D**) or human induced pluripotent stem cell (hiPSC)-derived islet organoids^10^ secrete insulin into the circulating co-culture medium. Secreted insulin was shown to stimulate glucose utilization in the liver model, composed of HepaRG hepatocytes and human hepatic stellate cells (HHSteC)^9, 11^ or hiPSC-derived liver organoids^10^. Simultaneously, as the glucose concentration in the co-culture medium fell from the initial hyperglycemic level to the normoglycemic range, insulin secretion subsided demonstrating a physiological feedback loop between the liver and the pancreas compartments. We have further shown that the capacity of HepaRG/HHSteC liver spheroids to utilize glucose from hyperglycemic co-culture medium decreased over time indicating the development of glucose dysregulation^9, 11^ (exemplified in **Fig. 1E**). This suggests that the pancreas-liver MPS could be used as a model to study the development of insulin resistance *in vitro*.

Here, we adapted this approach to develop a method for investigating diabetic glucose dysregulation on chip. We asked if our pancreas-liver MPS can represent an insulin resistance phenotype in the liver compartment and beta-cell adaptation/failure in the pancreas compartment. Due to the complex and dynamic nature of organ crosstalk, we combined the *in vitro* model with *in silico* modelling for hypothesis testing, data analysis, and informed decision-making. First, we resolved the elements responsible for the induction of glucose dysregulation. We investigated two medium supplements, glucose and glucocorticoid hydrocortisone (HCT), for their suspected influence on insulin sensitivity and beta-cell function. For this evaluation, we applied our recently developed mechanistic mathematical model^11^ to support the experimental design and to predict glucose and insulin responses in the pancreas-liver chip co-culture. By comparing computational predictions and experimental results, we identified HCT as a key factor inducing insulin resistance and beta-cell failure in the pancreas-liver MPS. Next, we demonstrated that our diseased two-organ model reflects several pathological alterations seen in patients suffering from glucocorticoid-induced diabetes. As we had observed signs of beta-cell adaptation in co-cultured islets, we hypothesized that these might be associated with factors secreted by the HepaRG/HHSteC liver spheroids that can induce beta-cell proliferation. Using combined transcriptome and proteome analysis, hit validation, and carrying out single-islet cultures, we showed that a liver spheroid-derived protein, IL-1R2, modulates islet proliferation. To test the repeatability, robustness, and transferability of the pancreas-liver MPS, and the inter-donor variability, we performed the MPS studies in two different laboratories using several different islet donors.

## RESULTS

### *In silico* supported experimental design

Previously, we suspected that the main driver for the glucose dysregulation observed in our pancreas-liver MPS^9, 11^ is the high glucose concentration (11 mM) of the co-culture medium because hyperglycemia is a known inducer of insulin resistance both *in vitro*^12^ and *in vivo*^13^. Therefore, we asked whether adapting the glycemic level to a normal blood glucose concentration (5.5 mM) could improve insulin sensitivity and, hence, maintain glucose utilization in the pancreas-liver model. However, when comparing normoglycemic and hyperglycemic conditions^11^ (**Fig. 1E**), we saw comparable glucose utilization during an *in vitro* adjusted glucose tolerance test (GTT)^9^ indicating that normoglycemia alone might not improve insulin resistance and glucose regulation. However, insulin concentrations during the GTT were lower in normoglycemia compared to hyperglycemia. Therefore, higher insulin resistance in the hyperglycemic condition might have been masked by a higher insulin secretion. This compensatory beta-cell adaptation might have led to comparable glucose utilization by the liver in the hyperglycemic and normoglycemic conditions. Based on these observations, we formed two hypotheses that could explain the development of insulin resistance in the pancreas-liver co-culture. The first hypothesis (H1; **Fig. 2A**, left graph) assumes that insulin resistance is caused by hyperglycemia alone, while the second hypothesis (H2; **Fig 2A**, right graph) assumes that insulin resistance is caused by a combination of hyperglycemia and an additional diabetogenic factor.

**Fig. 2.**
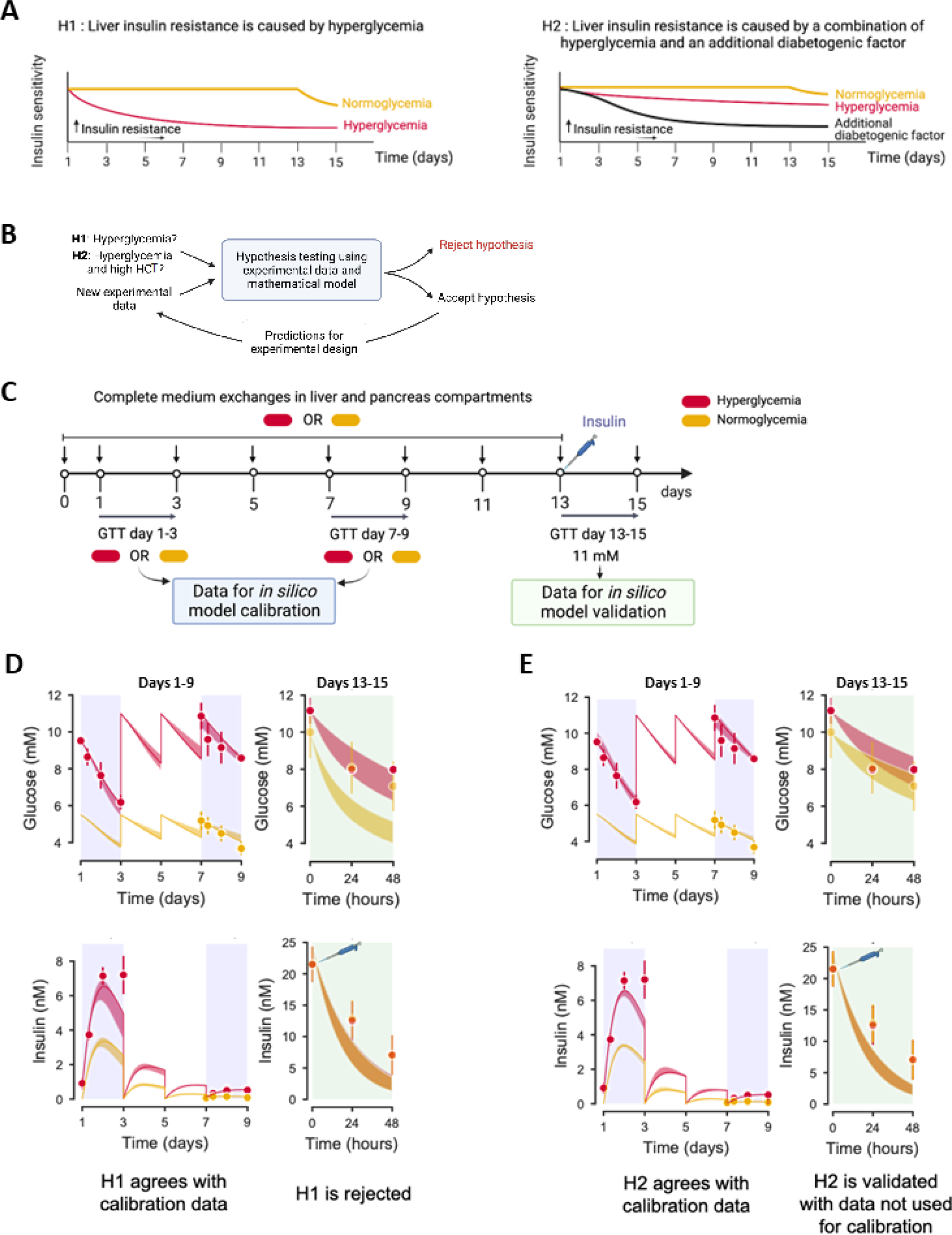
*In silico* supported experimental design for informed decision making. (**A, B**) Schematic representation of the hypothesis testing framework to unravel the cause of insulin resistance in the pancreas-liver MPS. (**A**) Here, we considered two competing hypotheses (H1, H2) for the development of insulin resistance in the pancreas-liver MPS. Hypothesis H1 (A, left graph) assumes that insulin resistance is caused by hyperglycemia alone, while hypothesis H2 (A, right graph) assumes that insulin resistance is caused by hyperglycemia in combination with an additional diabetogenic factor. **(B)** Hypothesis testing is an iterative process, where mathematical models constructed from experimental data are used to test and reject hypotheses. (**C**) *In silico* guided experimental design to test the proposed hypotheses. The computational model was first calibrated for donor-dependent variations using data from a glucose tolerance test (GTT) on days 1-3 and 7-9. Next, the computational model was used to select an insulin dose that was added to the co-cultures on day 13. This experimental design would lead to different glucose tolerances on day 13-15 and allow differentiation between H1 and H2. (**D,E**) Comparison between experimental data (dots) and model simulations (lines) in calibration phase (blue areas) and validation phase (green areas). The shaded areas (red and yellow) represent the model uncertainty. Both H1 and H2 agree with calibration data for glucose (top) and insulin (below) but only H2 predicts glucose response during the validation step on days 13-15. Data in panels D, E are presented as mean ± SEM, n= 3.

To study these hypotheses, we applied a computational hypothesis-testing approach (**Fig. 2B**; see Methods for details) using our recently described mathematical model of glucose and insulin interplay in the pancreas-liver co-culture^11^ (**Fig. S1A**). We designed a 15-day study involving sequential experimental and modelling iterations that would allow us to differentiate between the two hypotheses by spiking a defined amount of insulin to the co-culture medium on day 13 (**Fig. 2C).** The insulin dose is thereby selected based on the predictions of the mathematical model and would result in different glucose tolerance curves for hypothesis 1 and 2 (H1 and H2). To select the insulin dose, we first constructed mathematical models for both H1 and H2 (**Fig. S1B**). Next, we exposed the pancreas-liver co-cultures to either hyperglycemic or normoglycemic conditions for 13 days (**Fig. 2C**). Then, we calibrated the mathematical models for donor-dependent variations in the insulin secretion by feeding the models with recorded glucose and insulin concentrations at the beginning (GTT day 1-3) and in the middle (GTT day 7-9) of the co-culture study. Both H1 and H2 provided acceptable agreement with the experimental data (**Fig. 2D and E**, left graphs) according to a statistical c^2^ test (see Methods for details). Next, we used the calibrated mathematical models to select an insulin dose that, when spiked to the co-culture medium, would yield different predictions for glucose metabolism for hypotheses H1 and H2 (**Fig. S2**). When performing the GTT with the suggested insulin dose (24 nM), we saw similar utilization of an 11 mM glucose dose in both hyperglycemic and normoglycemic conditions (**Fig. 2D and E**, right graphs). Comparing the experimental data with the model predictions, we did not find a statistically acceptable agreement for H1 (**Fig. 2D**, right graphs) and therefore rejected the hypothesis that insulin resistance was induced by hyperglycemia alone. In contrast, H2 agreed with the experimental data (**Fig. 2E**, right graphs) according to c^2^ statistics. Therefore, we further investigated the hypothesis that insulin resistance was induced by a combination of hyperglycemia and an additional diabetogenic factor.

### Hydrocortisone and hyperglycemia drive insulin resistance on chip

As the computational hypothesis testing approach suggested that an additional diabetogenic factor is involved in the development of insulin resistance in the pancreas-liver co-culture, we suspected that an unphysiological HCT concentration may play a role. The standard HepaRG culture medium is supplemented with a high concentration of HCT^14^, a glucocorticoid that has an essential role in the differentiation and function of the liver^15^. However, glucocorticoids are known inducers of whole-body insulin resistance^16^ leading to a condition called glucocorticoid–induced or ‘steroid’ diabetes. In the liver, glucocorticoids have been reported to increase glucose production via gluconeogenesis^17^ and to promote hepatic lipid accumulation (steatosis)^18^ which is suspected to induce insulin resistance^19^. The HCT concentration in our original co-culture medium (50 µM)^9^ was several orders of magnitude higher than the free human plasma cortisol concentration (about 5.5-39 nM)^20^. Therefore, we hypothesized that the used HCT concentration in the co-culture medium might induce a diabetic phenotype in our pancreas-liver co-culture similar to that seen in patients suffering from steroid diabetes. Conversely, this means that reducing the HCT concentration to its physiological level might sustain insulin sensitivity by preventing the induction of gluconeogenesis and steatosis in the liver spheroids.

Glucocorticoids do not only affect the liver but have also been shown to impair the insulin secretion of beta cells *in vitro*^16, 21^ and *in vivo* in patients susceptible to beta-cell dysfunction^16, 21, 22^. Therefore, we first studied whether HCT has a direct negative effect on glucose-stimulated insulin secretion (GSIS) of the islets in our normoglycemic co-culture medium. We saw an inhibitory effect on insulin secretion already at 50 nM HCT (**Fig. 3A**) while a concentration of 5 nM HCT (near the physiological level) showed no difference to the untreated control. Next, we investigated whether a lower, physiological concentration of HCT in the co-culture medium would, first, maintain liver functions and improve insulin secretion in the pancreas-liver co-culture and, second, improve insulin sensitivity in the liver compartment, and the overall glucose regulation in the system. To study all variables, we maintained pancreas-liver co-cultures for two weeks in four different medium conditions using either high HCT (50 µM) or low HCT (10 nM) concentrations and either hyperglycemic or normoglycemic glucose concentrations (**Fig. 3B**).

**Fig. 3.**
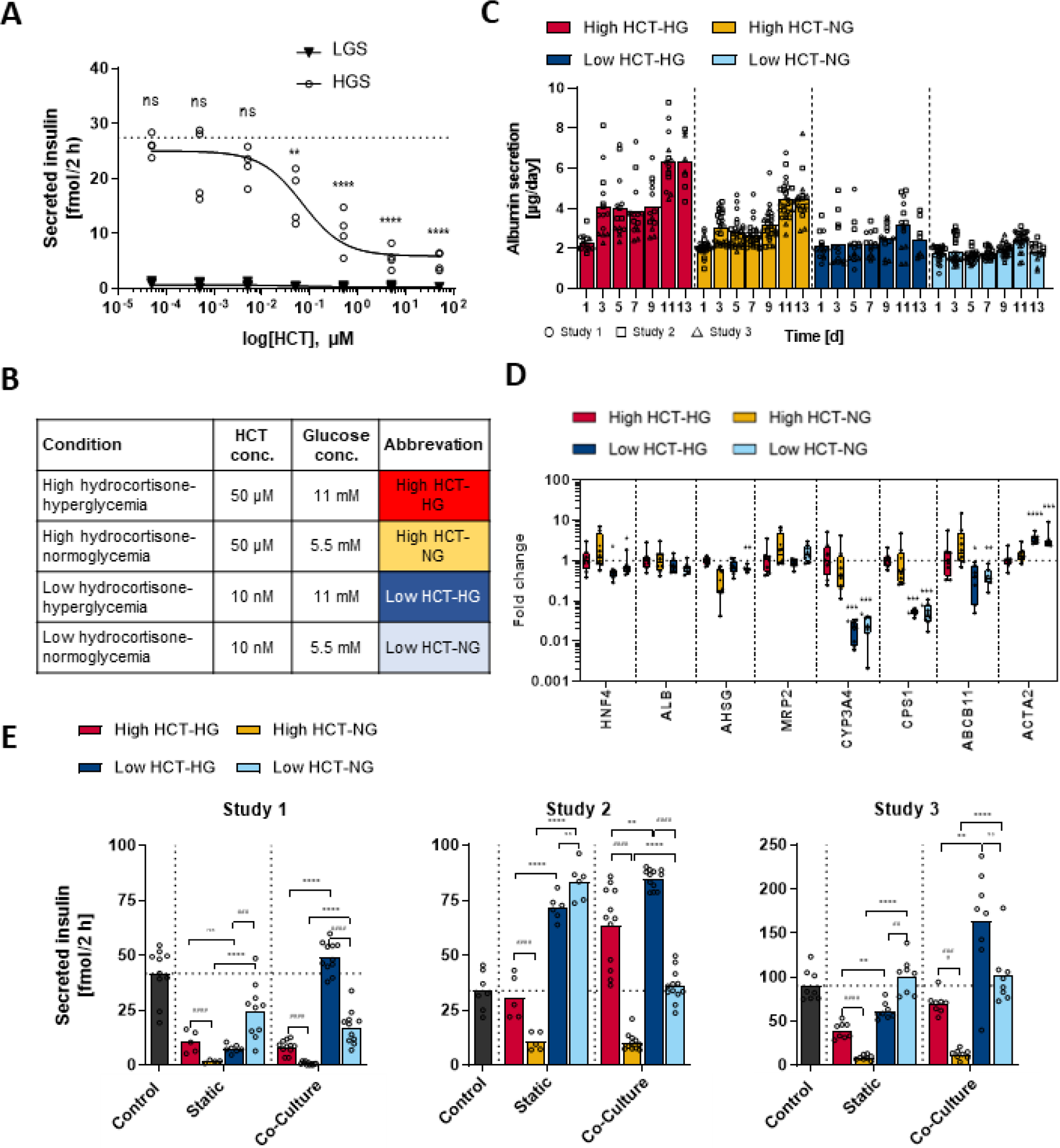
Hydrocortisone-induced insulin resistance and beta-cell dysfunction. (**A**) Hydrocortisone (HCT)-concentration dependent inhibition of glucose-stimulated insulin secretion (GSIS) of islets cultured in a normoglycemic co-culture medium. Model fitted to data using nonlinear regression. Differences to the control (no HCT, represented by dotted line) were evaluated by one-way ANOVA using Bonferroni’s multiple comparisons post-hoc test, **p < 0.01, ****p < 0.0001. LGS; Low-glucose stimulation with 2.8 mM glucose (triangles). HGS; High-glucose stimulation with 16.8 mM glucose (circles). (**B**) Overview of the four different experimental conditions studied in the pancreas-liver MPS. (**C**) Liver spheroid functionality shown by albumin secretion over the chip co-culture time. Symbols represent replicates from three independent studies. (**D**) Relative mRNA expression of key liver markers in the liver spheroids at the end of co-culture. Symbols represent replicates from two independent studies, total n = 8. Data shown as fold changes to the high HCT hyperglycemic condition. Differences between high and low HCT concentration for the same glucose concentration were evaluated by one-way ANOVA using Sidak’s multiple comparisons post-hoc test, * < 0.05, **p < 0.01, *** < 0.001, ****p < 0.0001. (**E**) GSIS response after high-glucose stimulation in islets cultured for 15 days in static or in chip co-culture. Data shown as a fold change to static islets cultured in the supplier’s maintenance medium that served as a control. Symbols represent individual islets. An individual donor was used for each study. Differences between selected pairs were evaluated by one-way ANOVA using Sidak’s multiple comparisons post-hoc test, asterisks (*) show comparison between high and low HCT concentrations for the same glucose concentration, hashtags (#) show comparison between hyper- and normoglycemia for the same HCT concentration, ns = not significant, **p < 0.01, ****p < 0.0001, ## < 0.01, ### < 0.001, #### < 0.0001. Data was log-transformed for normality.

First, we analysed the HepaRG/HHSteC liver spheroids’ functionality by following albumin secretion over time and measured mRNA expression of key markers of liver health. Here, we observed a stable albumin secretion at low HCT conditions while a high HCT concentration increased albumin secretion over time (**Fig. 3C**). This increase might be an initial sign of developing insulin resistance as patients with elevated serum albumin concentrations have an increased risk of developing T2D^23^. The expression levels of *HNF4A, ALB, AHSG, and MRP2* were not relevantly affected by lower HCT concentrations (**Fig. 3D**) while the expression of *CYP3A4* mRNA, a major drug-metabolism enzyme, *CPS1* mRNA, an enzyme participating in urea production, and *ABCB11* encoding BSEP, the major bile-acid transporter, were reduced at the low HCT conditions. This was not unexpected as glucocorticoids are known inducers of cytochrome P450 enzymes ^24^, the urea cycle (e.g., CPS)^25^, and hepatic bile acid transport^26^. Furthermore, the expression of *ACTA2* encoding alpha-smooth muscle actin was increased, suggesting the proliferation of HHSteCs when hydrocortisone concentration is reduced. This was also expected, as glucocorticoids are known for their anti-fibrotic effects^27^. In general, albumin secretion and the expression of liver-specific genes were preserved at the low HCT concentration, but some metabolic functions might be reduced compared to co-cultures maintained at high HCT concentrations.

Second, we studied the effect of different HCT and glucose concentrations on islets by analysing the GSIS after a dynamic co-culture or a static mono-culture. As demonstrated in three independent studies with individual islet donors, media with high HCT concentration resulted in a significant decrease in glucose-stimulated insulin secretion as compared to low HCT concentration in both hyper- and normoglycemia (**Fig. 3E**). Basal insulin secretion and stimulation index are reported in **Supplementary Fig. S3**. Interestingly, the hyperglycemic low HCT condition had a superior glucose-stimulated insulin secretion compared to the normoglycemic low HCT condition in all three co-culture studies but not in the corresponding static mono-culture studies. In addition, hyperglycemia increased insulin secretion in co-cultures exposed to high HCT concentration in studies 2 and 3, while this increase was not seen in the static mono-cultures. This effect is similar to the enhanced glucose-stimulated insulin secretion observed in healthy and prediabetic individuals as a response to a continuously rising blood glucose concentration^16, 21, 22^. Thus, our co-culture model might reflect this typical beta-cell adaptation mechanism. When comparing the three studies, we also found high variability in the donors’ ability to increase beta-cell function and, hence, adapt to developing insulin resistance. Indeed, islets from different individuals are known to have varying abilities for beta-cell adaptation^28^ as well as varying susceptibility for beta-cell failure through diabetogenic factors such as glucocorticoids^21^. In summary, the reduction of HCT concentration to a physiological level improved the glucose-stimulated insulin secretion showing that the effect of a high HCT concentration is in line with the beta-cell failure observed in patients suffering from steroid diabetes^22^.

To further evaluate whether a lower HCT concentration or normoglycemic glucose concentration, or these together, would lead to improved glucose regulation, beta-cell function, and insulin sensitivity during the co-culture, we performed a GTT in the pancreas-liver co-culture on day 1-3 (only hyperglycemic conditions) and on day 13-15 (all four conditions). To determine if the measured glucose and insulin responses could be explained by our hypothesis, we applied the following approach. First, we calibrated the computational model corresponding to hypothesis H2 using the experimental measurements from co-cultures exposed to high HCT (**Fig. 4A**). Then, we used the calibrated model to predict the expected insulin and glucose responses assuming that the lower HCT concentration would not affect insulin sensitivity, and the insulin secretion capacity would be maintained (**Fig. 4B**). By comparing these predictions to our experimental data, we found that the computational model can explain the measured responses indicating that low HCT concentration can indeed maintain the insulin sensitivity and beta-cell function in the pancreas-liver co-culture. This resulted in a maintained glucose tolerance as seen by stable glucose area under the curves (AUCs) over the culture time while these were increased at high HCT concentration (**Fig. 4C**). Confirming these findings, we saw similar responses in a repeated co-culture study with a difference that the glucose tolerance was only maintained in the low HCT-normoglycemic condition (**Fig. S4**).

**Fig. 4.**
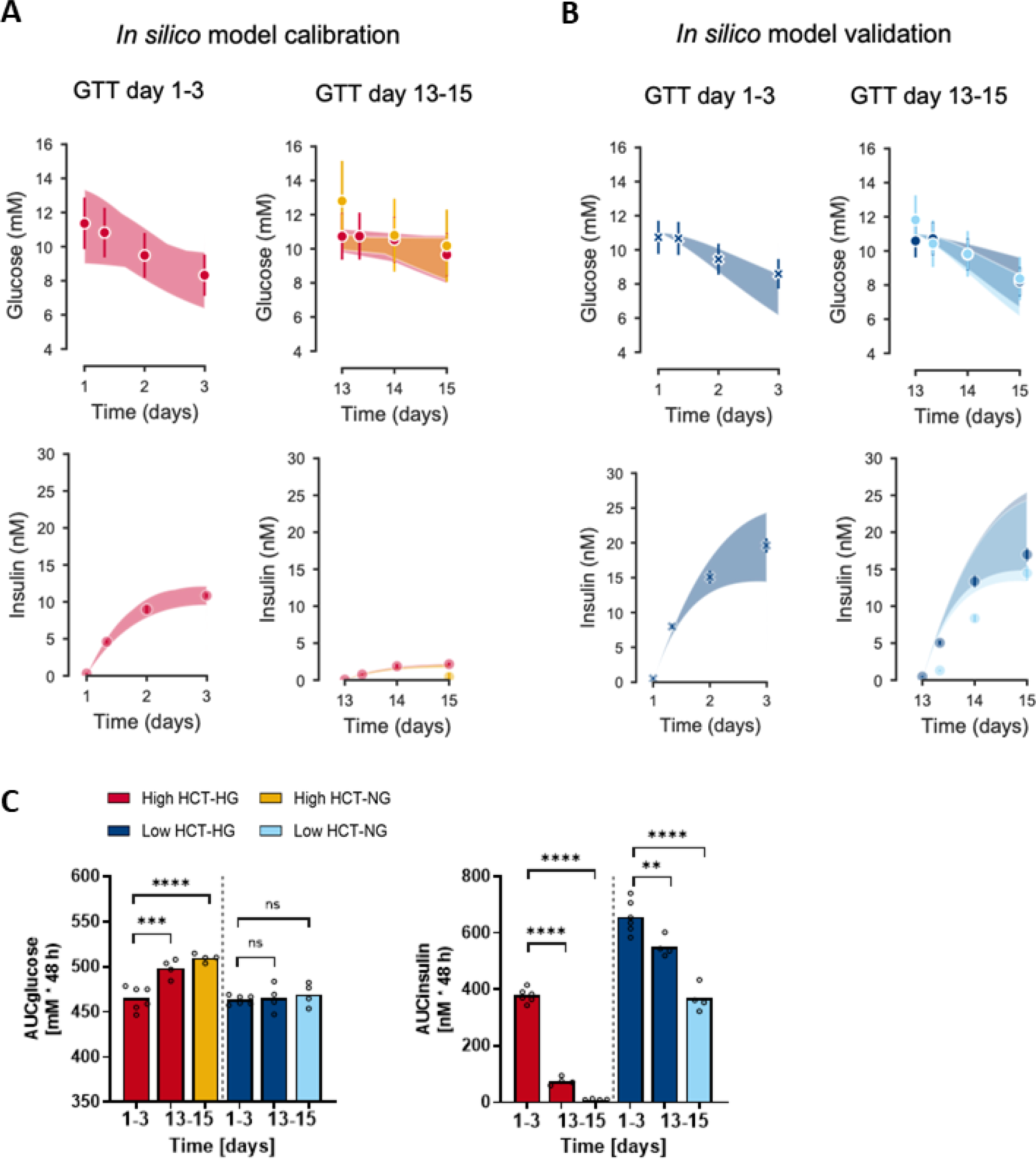
Glucose tolerance of the pancreas-liver co-culture. **(A)** The mathematical model was calibrated using experimental data measured in pancreas-liver co-cultures exposed to a high HCT concentration. Experimental measurements of glucose (dots, top row) and insulin (dots, bottom row) were acquired during GTTs on days 1-3 and 13-15 under normo- and hyperglycemic glucose concentrations. **(B)** The calibrated model was used to predict glucose (top row) and insulin responses (bottom row) at physiological HCT concentration. These predictions were compared to the corresponding experimental measurements (dots) from GTTs on day 1-3 (left) and 13-15 (right). Data points depicted with an X are used for baseline correction of insulin sensitivity and insulin secretion capacity. The shaded areas in panels a-b represent the model uncertainty, and the data are presented as mean ± SEM, n=4-6. (c) Area under the curve (AUC) for glucose (left) and insulin (right). Symbols represent individual pancreas-liver co-cultures from one study. Differences between selected pairs of conditions (day 13-15 compared to day 1-3) were evaluated by one-way ANOVA using Sidak’s multiple comparisons post-hoc test, ns = not significant, **p < 0.01, ***p < 0.001, ****p < 0.0001.

Altogether, we show that a ‘healthy’ pancreas-liver co-culture with stable liver function, beta-cell function, and glucose tolerance is achieved in a condition with low HCT concentration and normoglycemic glucose level. In contrast, a ‘diseased’ co-culture for representing impaired glucose tolerance accompanied by beta-cell dysfunction can be generated by using a high HCT-hyperglycemic medium. Therefore, we next focused on these two co-culture conditions as these are reflecting the healthy and diseased plasma concentrations of hydrocortisone and glucose observed *in vivo*. Data on the two intermediary conditions can be found in the supplementary material (**Figs. S5-S8**).

### Hepatic phenotype reflects glucocorticoid-induced diabetes

In patients with glucocorticoid-induced diabetes, hepatic insulin resistance is one factor contributing to dysbalanced glucose regulation and hyperglycemia^22^. Glucocorticoids increase endogenous glucose production by inducing the transcription of genes encoding gluconeogenic enzymes (e.g. glucose-6-phosphatase)^18, 29^. Moreover, glucocorticoids induce glycogen synthesis^30^ which increases the liver’s capacity to produce glucose. Furthermore, chronic elevation of glucocorticoid concentration has been linked to the development of a steatotic ‘fatty’ liver by increasing the gene transcription of several enzymes involved in *de novo* lipogenesis (including the fatty acid synthase)^18^. Excess fatty acids are partly converted to ketone bodies leading to elevated ketone levels in plasma^18, 31^.

To analyse how our liver model reflects the glucocorticoid-induced diabetic phenotype, we first looked at gene expression profiles of enzymes involved in glucose metabolism (**Fig. 5A**), ketogenesis (**Fig. 5B**), and lipid metabolism (**Fig. 5C**) in the co-cultured HepaRG/HHSteC spheroids. The diseased condition induced gene expression of glycogen synthase (*GYS2*) involved in glycogen synthesis, glucose-6-phosphatase (*G6PC*) involved in gluconeogenesis, HMG-CoA lyase (*HMGCL*) involved in ketogenesis, and fatty acid synthase (*FASN*) involved in *de novo* lipogenesis. Next, we confirmed these findings by performing separate analyses to evaluate glycogen storage, ketone body production, and lipid metabolism in the co-cultured HepaRG/HHSteC liver spheroids. Liver spheroids in the diseased co-cultures exhibited higher amounts of glycogen stores as shown by periodic acid-Schiff (PAS) staining (**Fig. 5D**), secreted 2.6-fold more 3-hydroxybutyrate (**Fig. 5E**), a diagnostic measure of diabetic ketoacidosis^32^, and accumulated more intracellular lipids as visualized by Nile Red staining (**Fig. 5F**) when comparing to spheroids in the healthy condition. These data indicate that the diseased liver model reflects several pathological alterations seen in patients suffering from steroid diabetes suggesting that the co-cultured HepaRG/HHSteC liver spheroids develop glucocorticoid-induced insulin resistance.

**Fig. 5.**
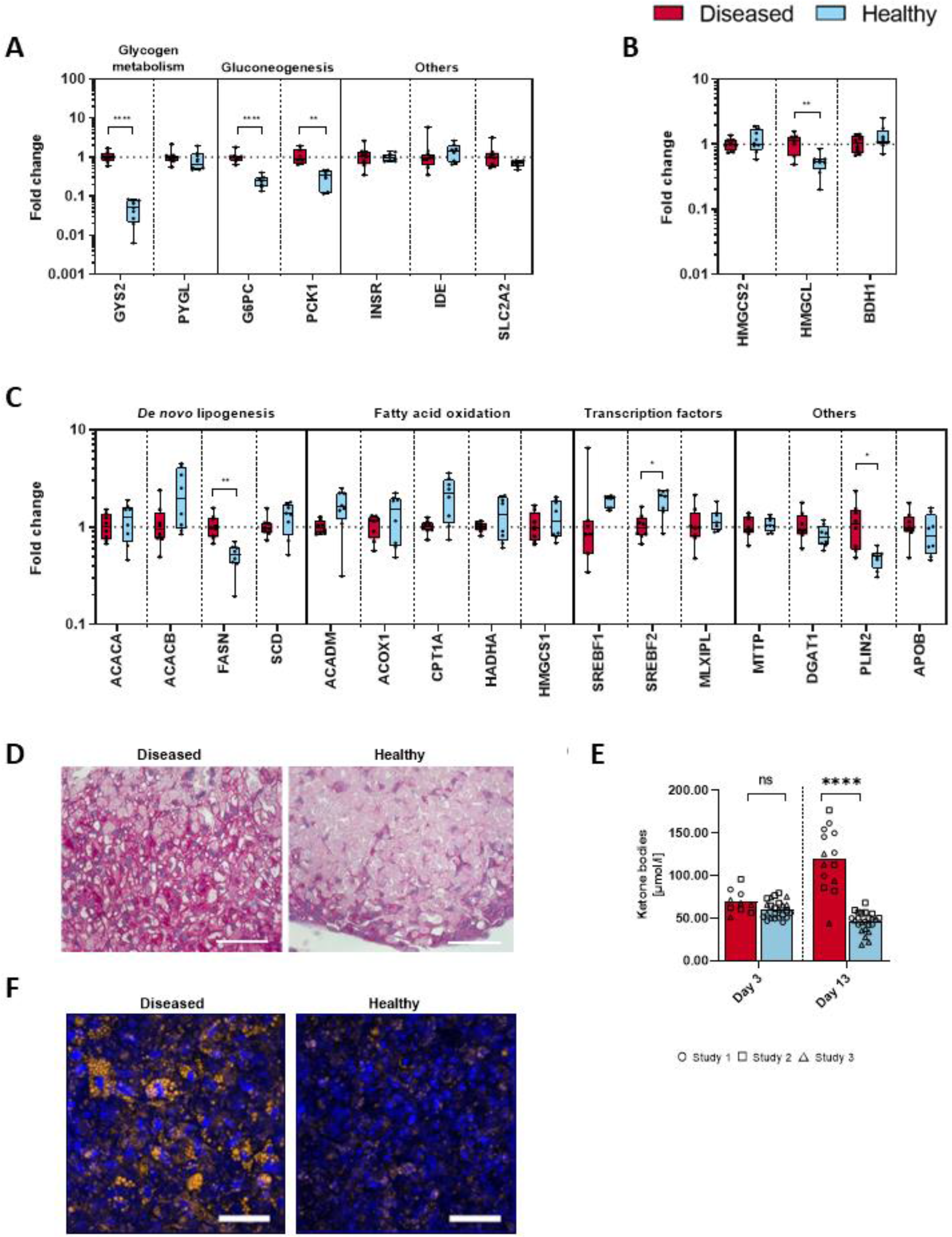
Insulin resistance-associated phenotype of HepaRG/HHSteC liver spheroids. **(A, B, C)** Gene expression of enzymes involved in hepatic glucose metabolism **(A),** ketone-body synthesis (**B**), and lipid metabolism (**C**). Data shown as fold change between diseased (11 mM glucose, 50 µM HCT) and healthy conditions (5.5 mM glucose, 10 nM HCT) in a box-whisker plot with geometric mean and min-max values. Symbols represent replicates from two independent studies, total n = 8. Differences between the diseased and the healthy condition were evaluated by multiple t-tests using the Holm-Sidak method for multiple comparisons without assuming a consistent standard deviation. (**D**) Glycogen storage visualized by periodic acid-Schiff (PAS) staining. Scale bar, 50 µm. (**E**) Ketone-body synthesis represented by 3-hydroxybutyrate concentration in the co-culture supernatants. Symbols represent co-culture replicates from three independent studies. Differences between day 3 and day 13 were evaluated by multiple t-tests using the Holm-Sidak method for multiple comparisons without assuming a consistent standard deviation. (**F**) Intracellular lipid vesicles visualized by Nile Red staining (amber colour). Blue denotes DAPI-stained nuclei. Scale bar, 50 µm.

### Evaluation of liver-derived effects on islet functions

Individuals with insulin resistance do not necessarily develop glucose dysregulation and diabetes as beta-cells can compensate for the increased insulin demand by either increasing in number (proliferation or transdifferentiation) or enhancing their secretory output, or both^33^. Previously, several studies have demonstrated that organs, including the liver, secrete proteins into the bloodstream which stimulate insulin secretion and proliferation of islets^34^. To evaluate whether the observed improvement in insulin secretion in the co-culture as compared to the static mono-culture (**Fig. 3E**) could be explained by an increased islet cell number, we developed a robust cell proliferation assay using 5-ethynyl-2’-deoxyuridine (EdU) incorporation, automated high-throughput confocal microscope imaging, and automated image analysis (**Fig. S9**). When the islets were cultured in the disease condition, proliferation did not differ between the chip co-culture and static mono-culture (**Fig. 6A**) suggesting that other beta-cell adaptation mechanisms than increased cell mass contribute to the improved insulin secretion capacity seen in co-cultures (**Fig. 3E**). Instead, in the healthy condition proliferation was significantly increased in co-cultured islets as compared to the mono-cultured islets.

**Fig. 6.**
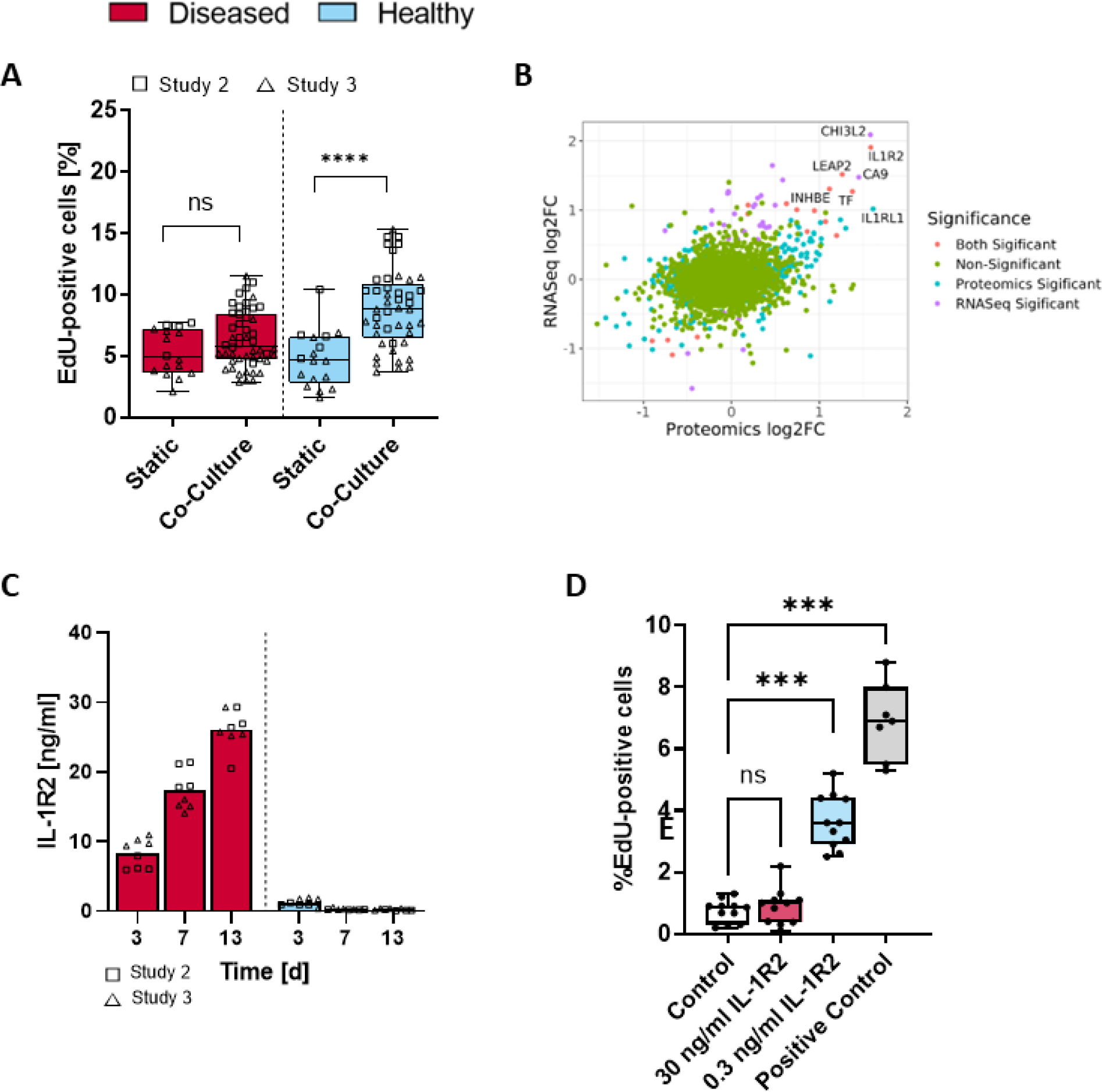
Evaluation of liver-derived effects on islet functions. (**A**) Proliferation of islets in the diseased (11 mM glucose, 50 µM HCT) and healthy (5.5 mM glucose, 10 nM HCT) conditions at the end of the culture as analysed by the percentage EdU-positive cells. Squares (study 2) and triangles (study 3) represent individual islets from two independent co-culture studies. Differences between the static and the co-culture were evaluated by a two-tailed unpaired t-test, ns = not significant, ****p < 0.0001. (**B**) Multi-omics analysis on the effect of hyperglycemia vs. normoglycemia on liver-secreted proteins. Data from RNASeq (HepaRG/HHSteC spheroids) and proteomics (co-culture supernatants) were merged at the gene level. Point colour indicate significance of change (FDR<0.05 and p<0.05 for RNASeq and proteomics data, respectively). Gene names are marked for genes with RNASeq and proteomic log2 fold-changes > 1. Transcriptomics data is from three independent studies and proteomics data from four independent studies. (**C**) Secretion of IL-1R2 in the co-cultures over time. Squares (study 2) and triangles (study 3) represent individual islets from two independent co-culture studies, total n = 8. (**D**) IL-1R2 stimulates cell proliferation at low dose (0.3 ng/mL) but not at the high dose (30 ng/mL) in islets mono-cultured in static condition in Human Islet Maintenance Medium (InSphero). Control, 0 ng/mL IL-1R2. Positive control, low hydrocortisone-normoglycemic (healthy) co-culture medium. Symbols represent individual islets. Differences to the control were evaluated by one-way ANOVA using Dunnett’s multiple comparisons post-hoc test, ns = not significant, ***p < 0.001.

Therefore, we performed exploratory transcriptome and proteome analyses evaluating the influences of hyperglycemia and normoglycemia on liver-secreted proteins in the chip co-cultures. When combining RNA sequence analysis of co-cultured HepaRG/HHSteC liver spheroids and proteomics analysis of supernatants at the end of the chip co-culture, IL-1R2 was the most upregulated protein in the hyperglycemic condition (**Fig. 6B**). IL-1R2 is a decoy receptor for IL-1beta which is an inflammatory cytokine associated with diabetes and especially beta-cell dysfunction^35^, thus a target for diabetes therapies. In *in vitro* studies, animal models, and clinical trials, inhibition of interleukin-1 receptor (IL-1r) has been shown to enhance beta-cell survival and function^36–39^. Therefore, we hypothesized that IL-1R2 could have a similar effect on islets by reducing the detrimental free IL-1beta concentration. To test this, we first quantified secreted IL-1R2 in the chip co-cultures over time and noticed a significant upregulation in IL-1R2 secretion in the disease condition (**Fig. 6C**). To confirm that IL-1R2 is solely produced by the liver compartment, we analysed IL-1R2 secretion in static mono-cultured islets and saw no secretion (**Fig. S10A**). Next, we treated islets in static mono-culture with 30 ng/mL or 0.3 ng/mL of IL-1R2 mimicking the measured levels in the diseased and healthy condition, respectively. Compared to untreated control, we observed a 4.9-fold increase in proliferation measured as a proportion of EdU-positive cells in islets treated with 0.3 ng/mL of IL-1R2 (**Fig. 6D**). In contrast, 30 ng/mL of IL-1R2 did not affect proliferation. These results suggest that liver-derived IL-1R2 may be one factor impacting islet proliferation in the healthy pancreas-liver co-culture (**Fig. 6A**). Interestingly, it has been reported that low, but not high, IL-1beta concentration has beneficial effects on islet functionality^40^. Therefore, we hypothesize that the low IL-1R2 concentration in the healthy condition might have reduced the IL-1beta concentration to a beneficial range while the high IL-1R2 concentrations resulted in ineffectively low IL-1beta concentrations. In line with an earlier observation that proliferating beta cells have an impaired insulin response^41^, we observed reduced glucose-stimulated insulin secretion at low IL-1R2 concentration (**Fig. S10B**).

## DISCUSSION

Preclinical T2D studies rely on animal models because the standard *in vitro* single-cell or single-organ cultures cannot replicate organ-to-organ crosstalk essential for multisystem disorders. However, the animal models are genetically and physiologically different from humans^4^, and more accurate human-based preclinical models are therefore needed. Here, we describe a diseased pancreas-liver MPS model that can replicate hallmark features of diabetic dysregulation both in the liver and pancreas compartments.

We applied computational modelling to guide hypothesis testing, experimental design, and data interpretation and showed that the pancreas-liver MPS exhibits a diabetic phenotype including glucose dysregulation, insulin resistance, and beta-cell dysfunction when the chips are exposed to a medium reflecting diabetic glucose and glucocorticoid concentrations. This experimental-computational hybrid approach is important for the correct interpretation of multi-organ MPS data as cross-organ feedback loops are hard or even impossible to unravel by pure reasoning. Notably, computational modelling also allows *in vitro*–to–*in vivo* translation. We recently showed that pancreas-liver MPS results can be translated to humans by using mechanistic mathematical modelling even if some of the MPS characteristics do not reflect human physiology, such as cell-to-liquid ratio and the flow rate, since these can be corrected in the mathematical models^11^.

By using the described pancreas-liver MPS, we demonstrated that the diseased condition with hyperglycemic glucose level and high hydrocortisone concentration reflected several pathological alterations seen in patients suffering from glucocorticoid-induced diabetes. In the liver compartment, these included steatosis, diminished glucose utilization as well as increased ketone-body production, and beta-cell dysfunction in the islet model. We evaluated the model in two laboratories and observed low inter-experimental and inter-laboratory variation. We also showed that the method is effective in three pancreatic islet donors. Therefore, the pancreas-liver model offers a human-based system to study diabetic glucose dysregulation as an alternative to animal models. Importantly, the inter-donor comparison between different islet productions allows the investigation of varying susceptibilities for beta-cell damage by diabetogenic factors as well as their varying ability for beta-cell adaptation which is not possible in animal models due to their monogenetic background. To accommodate studies on inter-donor variability also for the liver part, we are currently developing a pancreas-liver MPS method with primary human hepatocytes. Additionally, the model could be used to study long-term drug exposures, for example glucocorticoids, as such studies are not feasible in human volunteers due to the risk of irreversible negative effects^21^.

Rodent islets are known to have higher beta-cell adaptation capacity via proliferation as compared to humans^42, 43^ and, thus, they are not an ideal model for finding human-relevant targets. Having observed that islets cultured in the pancreas-liver model have enhanced proliferation as compared to the islet in static cultures, we explored liver-derived proteins that might be responsible for the stimulation. We showed that IL-1R2, secreted from the liver compartment, can modulate islet proliferation. These findings did not only confirm that the liver and pancreas compartments exhibit disease-relevant crosstalk on-chip but also further amplifies that the described multi-organ model can be used to study new targets and therapies for diabetic patients.

Our current multi-organ MPS would benefit from having another target tissue for insulin action such as an adipose-tissue model. While the liver plays a central role in controlling the glucose metabolism, adipose tissue modulates glucose and lipid metabolism via releasing free fatty acids, adipokines, and proinflammatory cytokines^44^. Patients on corticosteroids have reduced glucose uptake and increased lipolysis in the adipose tissue leading to both elevated glucose and fatty-acid levels in plasma^22^. However, translational *in vitro* models of adipose tissue are not trivial to establish^45^, especially because the adipose tissue is highly heterogeneous^46^. Recently, Slaughter *et al*. successfully coupled liver and adipose models on chip with functional adipokine signalling for 14 days^47^. Pancreas-liver-adipose MPS would reflect insulin resistance pathophysiology more broadly and allow investigations of emerging therapies targeting adipose tissue^48^. Furthermore, the use of hiPSC-derived organ models could reflect the highly heterogenous disease progression and allow the testing of treatment options on a patient-derived diabetes model on-a-chip.

Together, the pancreas-liver *in vitro* and *in silico* hybrid model for glucose dysregulation enables diabetes research in a human-based preclinical system. A partnership of advanced cell models and computing is a necessity for studies on multisystem diseases with complex organ-to-organ communication. The model should facilitate drug discovery by serving as a platform for studies on disease mechanisms, target identification, and candidate drug evaluation.

## MATERIALS AND METHODS

### Liver spheroid formation

All cell cultures were maintained at 37 °C and 5% CO_2_ and conducted according to good cell culture practice^49^. We used terminally differentiated human HepaRG cells as a hepatocyte model as their gene expression profiles, regulatory pathways, and functional glucose machinery and lipid metabolism are similar to that in primary human hepatocytes^50–52^. Furthermore, a functional insulin responsiveness was described for HepaRG cells^51^ which is further improved in a three-dimensional spheroid culture^9^. Before liver spheroid formation, differentiated HepaRG hepatocyte-like cells (HPR116080, Biopredic, Lot HPR116NS080003 and HPR116239-TA08 or NSHPRG, Lonza, Lot HNS1014) were pre-cultured as previously described with a modification to medium composition^9^. Glucose and insulin concentration of the pre-culture medium were adjusted to physiological levels resulting in the following composition: Williams’ medium E (P04-29050S4, PAN-Biotech, w/o glucose, w/o L-glutamine, w/o phenol red) supplemented with 10% foetal bovine serum (FBS; 35-079-CV, Corning or 10270-106, Gibco), 5.5 mM glucose (25-037-CIR, Corning or 072397, Fresenius Kabi), 1 nM insulin (P07-4300, PAN-Biotech or 12585-014, Gibco), 2 mM GlutaMax (35050-061, Gibco), 50 µM hydrocortisone hemisuccinate (H4881, VWR or H2270, Sigma Aldrich), 50 µg/ml gentamycin sulphate (30-005-CR, Corning or 15710-049, Gibco) and 0.25 µg/mL amphotericin B (30-003-CF, Corning).

Primary human hepatic stellate cells (HHSteC, S00354, BioIVT, Lot PFP) were expanded in Stellate Cell Medium (5301, ScienCell) supplemented with Stellate Cell Growth Supplement, 2% FBS and 1% penicillin/streptomycin, and cryopreserved in FBS with 10% DMSO (23500.297, VWR). The HHSteCs (p3-4) were thawed at least two days before spheroid formation and pre-cultured in stellate cell medium until spheroid formation.

Liver spheroids were formed for 3 days in 384-well spheroid microplates (3830, Corning) with 24,000 differentiated HepaRG hepatocytes and 1,000 HHSteCs per spheroid as described previously^9^. Once compact spheroids had formed, 40 spheroids were collected into a 24-well ultra-low attachment plate (3473, Corning) for each co-culture replicate, and incubated in 1 mL pre-culture medium overnight on a 3D rotator (PS-M3D; Grant-bio) before transfer to the islet-liver co-culture.

### Pre-culture of pancreatic islets

We used commercially available human pancreatic islet microtissues (MT-04-002-0, InSphero) as a pancreatic islet model. The microtissues are manufactured from a dissociated human pancreatic islet suspension and have a defined cell number. After arrival, the pancreatic islet microtissues (termed islets throughout the manuscript) were maintained for 5 days in Akura™ 96 Spheroid Microplate (CS-09-004-01, InSphero) according to the manufacturer’s instructions. Medium was exchanged every 2-3 days with 70 µL of Human Islet Maintenance Medium (CS-07-005-02; InSphero). Donor for the hypothesis testing study was male, 52 years, with BMI of 29.6 and HbA1c of 5.4%. Donor for study 1 was male, 29 years, with BMI of 22.2 and HbA1c of 5.5%. Donor for study 2 was male, 26 years, with BMI of 24.1 and HbA1c of 5.1%. Donor for study 3 was male, 55 years, with BMI of 30.9 and HbA1c of 5.6%.

### Pancreas-liver chip co-culture

We performed co-cultures with islets and liver spheroids on a commercially available multi-organ-chip Chip2 (TissUse) platform (**Fig. 1D**). This MPS has two culture compartments for the integration of spatially separated organ models. The culture compartments are interconnected by a microfluidic channel. An on-chip micropump drives a pulsatile flow supporting long-term perfusion and communication between the organ models. Design and fabrication of the Chip2 were described previously (Schimek, 2013, Wagner, 2013).

Three days before insertion of the organ models, the chips were prepared for cultivation by replacing the storage buffer with 300 µL co-culture medium in each culture compartment (total volume per circulation was 605 µL). The chips were connected via air tubes to the control unit (HUMIMIC Starter) operating the on-chip micropump. The control unit was set to 0.45 Hz, 500 mbar pressure and –500 mbar vacuum resulting in an average volumetric flow rate of 4.94 µL/min between the culture compartments.

On the day of organ model transfer, the liver spheroids were washed twice with PBS to remove insulin from pre-culture medium. Subsequently, the liver spheroids were equilibrated for at least 2 hours in an insulin-free co-culture medium composing of Williams’ medium E (w/o glucose, w/o L-glutamine, w/o phenol red), 10% FBS, 2 mM GlutaMax, 50 µg/mL gentamycin sulphate, and 0.25 µg/mL amphotericin B. Glucose concentration was 5.5 mM in the normoglycemic condition and 11 mM in the hyperglycemic condition, and hydrocortisone concentration was either 10 nM or 50 µM (indicated in each study and condition). The islets were similarly equilibrated in the co-culture medium for at least 2 hours. After equilibration, 40 liver spheroids and 10 pancreatic islets were transferred to their respective culture compartment with 300 µL of fresh co-culture medium. Liver spheroids were collected using a wide-bore filter tip (T-205-WB-C-R-S, Corning), and carefully transferred into the liver compartment. In parallel, 10 islets were collected into a 1.5 mL microtube and pelleted by a brief centrifugation (1 min, 200 g) and transferred to the pancreas compartment. Alternatively, the islets were collected using an electronic single-channel pipette (Xplorer plus, Eppendorf) and directly transferred into the chips. The co-cultures were dynamically incubated at 37 °C and 5% CO_2_ using the same settings in the control unit as described above. The co-culture medium in both culture compartments was exchanged completely after 24 hours (adaptation time to the dynamic culture) and subsequently every 48 hours for a total co-culture duration of 15 days.

In studies 1, 2, and 3, some islets were statically cultured in parallel to the chip co-cultures. The islets were kept in Akura™ 96 Spheroid Microplate and medium was exchanged according to the co-culture study design.

### IL-1R2 treatment

Islets in Akura™ 96 Spheroid Microplate were treated with 0.3 ng/mL or 30 ng/mL of human recombinant IL-1R2 protein (10111-H08H, Sino Biological) for 16 days. Medium with IL-1R2 was renewed three times a week. For the last five days of culture, medium was also supplemented with 10 µM EdU for proliferation analysis. Islets cultured in Human Islet Maintenance Medium served as an untreated control and islets cultured in insulin-free co-culture medium with 11 mM glucose and 10 nM hydrocortisone served as a positive control in the proliferation assay. After finishing the culture, the islets were analysed for their glucose-stimulated insulin secretion and proliferation using EdU incorporation assay as described below. The islet donor was a female, 32 years with BMI of 25.6 and HbA1c of 5.1%.

### Glucose tolerance test

We performed GTT as described previously^9^ at different timepoints during the co-culture. Briefly, we exchanged the co-culture medium in both culture compartments with a co-culture medium containing 11 mM glucose (−300 µL, +315 µL) and collected 15 µL of supernatant samples at 0, 8, 24, and 48 h to monitor glucose and insulin concentrations. To obtain sufficient sample volumes for the analysis, samples from the liver and pancreas compartments were pooled. For optimal sample recovery, samples were stored in 96-well PCR plates (30133358, Eppendorf) and sealed using aluminium foil to minimize evaporation during storage. Samples were stored at −80 °C until glucose and insulin measurements.

### Glucose-stimulated insulin secretion

To assess functionality of the islets after the co-culture, we extracted islets from the chips, transferred into Akura™ 96 Spheroid Microplate, and performed GSIS on individual islets. The islets were first washed twice with 70 µL of Krebs-Ringer solution containing 2.8 mM glucose (low glucose solution), followed by equilibration in 70 µL of low glucose solution for 1-2 hours. Next, the islets were washed twice with 70 µL of low glucose solution and incubated for 2 hours in 50 µL of low glucose solution to measure basal insulin secretion. Following this, the islets were washed once with 70 µL of Krebs-Ringer solution containing 16.8 mM glucose (high glucose solution) and subsequently incubated in 50 µL of high glucose solution for 2 hours to measure the glucose-stimulated insulin secretion. Basal and glucose-stimulated samples were collected after incubations and stored at −80 °C until insulin measurement.

### Analysis of soluble markers

Supernatants of cell culture medium collected during medium exchanges were analysed for albumin (10242, Diagnostic Systems) and 3-beta-hydroxybutyrate (Autokit 3-HB, Fujifilm Wako) on an Indiko Plus chemical analyzer (Thermo Fisher Scientific) according to the manufacturer’s instructions. IL-1R2 concentrations were determined in culture supernatants by an ELISA assay (EHIL1R2, Thermo Scientific) according to the manufacturer’s instructions. Samples taken during the GTT were analysed for glucose (1070-500, Stanbio Laboratory) and insulin (10-1113-10, Mercodia) according to the manufacturer’s instructions.

### Computational models

#### Hypothesis testing using computational modelling

We used mathematical modelling as a tool to test mechanistic hypotheses on experimental data. A mechanistic hypothesis corresponds to a formulation of causal mechanisms key to produce the observed behaviour in the data. Hypothesis testing via mathematical modelling is an iterative approach (**Fig. 2B**). In the first step, the existing hypotheses are translated into a set of mathematical equations (i.e., corresponding mathematical models). We considered two hypotheses for the observed glucose and insulin responses in the pancreas-liver MPS: H1, “Insulin resistance is caused by hyperglycemia alone”, and H2, “Insulin resistance is caused by a combination of hyperglycemia and an additional diabetogenic factor”.

The second step involves the acquisition of experimental data and fitting the mathematical models to these data by optimization of the model parameters. The hypotheses are initially evaluated based on the outcome of this optimization. If the mathematical model cannot provide an acceptable agreement with the data, according to statistical analyses, then the corresponding hypothesis is rejected and must be revised. On the other hand, if the model can provide an acceptable agreement with data, the corresponding hypothesis is not rejected. The non-rejected models can then be used to generate uniquely identified predictions with uncertainty^53^, that allow for designing new experiments that could distinguish between the remaining hypotheses. The experiments are performed, and the predictions are compared against the new experimental data. If the model predictions agree with the experimental data, the corresponding hypothesis is accepted. On the contrary, if the predictions do not agree with the experimental data, the model is rejected and a new iteration in the hypothesis testing cycle is performed. Several iterations can be performed until a final model has been found.

In the following, we describe the modelling process, the mathematical model with its equations, and the hypothesis testing procedure.

#### A computational model for glucose metabolism in the pancreas-liver co-culture

We used our previously developed computational model^11^ as a basis to implement the hypotheses studied in this paper. The model describes glucose metabolism in the pancreas-liver co-culture (**Fig. S2B**). More specifically, it describes crucial biological processes underlying glucose regulation on a short-term basis (meal response), as well as long-term changes in physiological variables related to impaired glucose homeostasis, such as insulin resistance and beta cell adaptation. This model was constructed based on experimental data from seven independent studies corresponding to seven different islet donors^11^.

The computational model is formulated using ordinary differential equations (ODEs), which have the following general structure:

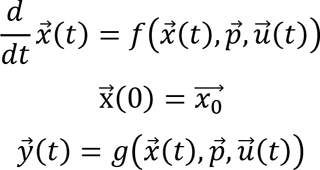

where 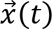 is the state vector describing the dynamics of concentrations or amounts and 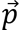 are the parameters, which here correspond to kinetic rate constants. 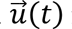 is a vector containing the external inputs. 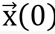 contains the initial conditions, i.e., the values of the states at 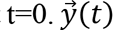 are the simulated model outputs, which correspond to the measured experimental signals. *f* and *g* are non-linear smooth functions that describe a set of mechanistic assumptions.

#### Derivation of the computational model

The computational model is based on the interplay between two components corresponding to different time scales: fast (hours) and slow (weeks). The fast model describes glucose and insulin dynamics between medium exchanges, which take place every 48 hours. The slow model describes the dynamics of long-term variables representing disease progression, such as the development of insulin resistance in the liver spheroids and beta-cell adaptation in the islets. The interplay between these two models allows short-term variables to impact long-term disease progression (e.g., impact of daily glucose levels on insulin resistance and beta cell volume) and vice versa. The model includes two compartments, each of them representing a specific culture compartment in the MPS (liver or pancreas) comprising a corresponding organoid and co-culture medium. The compartments are connected in a closed loop, with circulating medium determined by a flow rate parameter. The model equations are described in detail previously^11^ and summarized below.

Glucose content in the co-culture medium within the liver compartment varies with glucose dosing to the system, endogenous glucose production and glucose uptake by the liver spheroids, as well as glucose inflow from and outflow to the pancreas compartment:

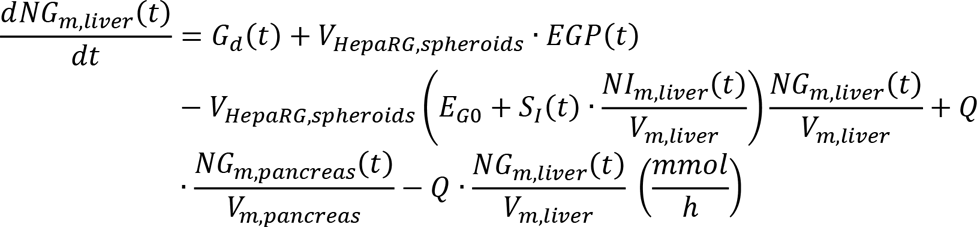

where *NG*_*m,liver*_(*t*) and *NG*_*m,pancreas*_(*t*) are the number of glucose molecules (mmol) in the culture medium corresponding to the liver and pancreas compartments, respectively, and *NI*_*m,liver*_(*t*) is the number of insulin molecules in the co-culture medium within the liver compartment (mIU). The glucose input rate *G*_*d*_(*t*) (mmol/h) defines glucose variations due to media exchanges, and *EGP*(*t*) describes endogenous glucose production in the liver spheroids (mmol/L/h). *EGP*(*t*) was set to zero based on the observed decline in glucose levels below normoglycemia (5.5 mM) in our system. Glucose uptake by the liver spheroids accounts for both insulin-independent uptake, determined insulin-independent glucose disposal rate *E*_*G*0_(1/h), and an insulin-dependent uptake regulated by the insulin sensitivity of the liver spheroids *S*_*I*_(*t*) (L/mIU/h). The parameters describing the flow rate between culture compartments (*Q* (L/h)), the total volume of HepaRG cells in the liver spheroids (*V*_*HepaRG,spheroids*_(L)) and the volume of co-culture medium in the liver and pancreas compartments (*V*_*m,liver*_and *V*_*m,pancreas*_(L), respectively) account for the operating conditions in the MPS.

In the computational model, insulin sensitivity of the liver spheroids *S*_*I*_(*t*) decreases progressively from its initial value at the beginning of the co-culture *S*_*I*0_ (L/mIU/h), as the liver spheroids are exposed to hyperglycemic concentrations (i.e. above normoglycemia) over time. This decrease is determined by the maximal fractional reduction *I*_*max,Si*_, and with half of the maximal fractional reduction occurring at *EC*50_*Si*_(mmol·h/L). The hyperglycemic periods are quantified by the integral of excess glucose *G*_*int*_(*t*):

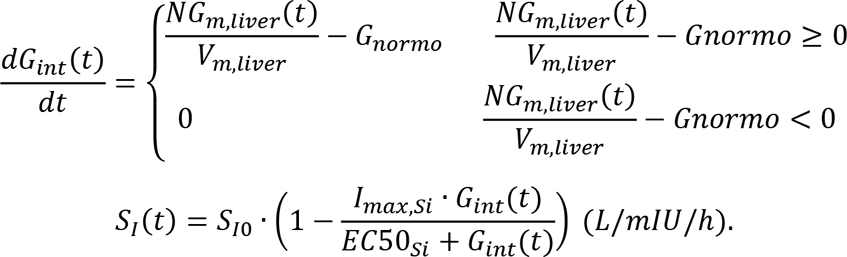

Glucose content in the pancreas compartment is described as:

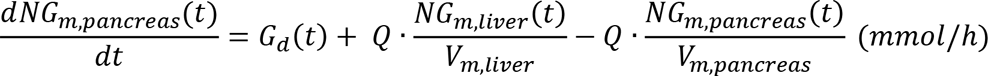

Insulin content in the pancreas compartment depends on the release of insulin from beta cells in the islets, and insulin inflow from and outflow to the liver compartment. Insulin release from the beta cells was modelled as a combination of the volume of beta cells in the islets (*V*_*β,islets*_(*t*) (L)), the insulin secretion capacity per unit volume of beta cells (denoted σ(*t*) (mIU/L/h)), and the glucose concentration resulting in half-of-maximum response to insulin (denoted *EC*50_*I*_(mmol/L)). The full equation describing insulin content in the pancreas compartment then becomes:

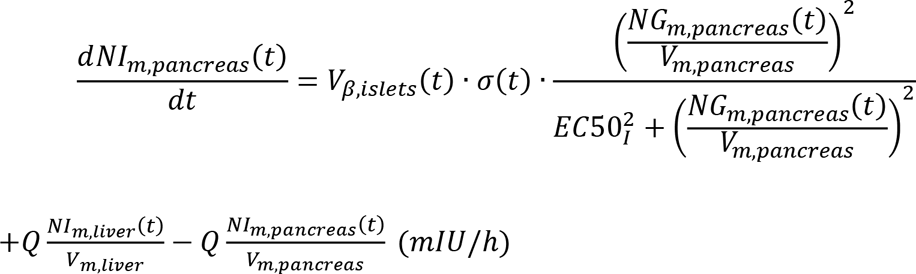

where *NI*_*m,pancreas*_(*t*) and *NI*_*m,liver*_(*t*) are the number of insulin molecules (mIU) in the pancreas and the liver compartment, respectively.

Furthermore, the insulin secretion capacity of the b cells was modelled as a decreasing function of time, determined by the parameter *α* (h^2^):

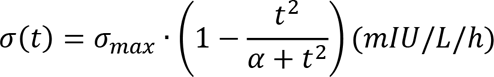

where σ_*max*_ (mIU/L/h) represents the maximal insulin secretion rate of the beta cells (i.e. at the beginning of the co-culture).

The variable *V*_*β,islets*_(*t*) (L) describes the changes in volume of beta cells in the pancreatic islets over the co-culture time, according to the following equation:

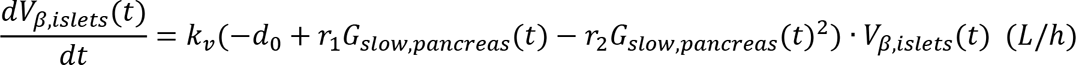

where *d*_0_is the death rate at zero glucose (h^-1^), *r*_1_ = *r*_1,*r*_ + *r*_1,*a*_ (L/mmol/h) and *r*_2_ = *r*_2,*r*_ + *r*_2,*a*_ (L^2^/mmol^2^/h), where *r*_1,*r*_, *r*_1,*a*_ (L/mmol/h), *r*_2,*r*_, *r*_2,*a*_ (L^2^/mmol^2^/h) are parameters that determine the dependence on glucose of the replication and apoptosis rates. The parameter *k*_*v*_ was introduced to account for potential differences in behaviour between islets in our *in vitro* system and rodent islets in the model of Topp et al.^54^.

The variable *G*_*slow,pancreas*_(*t*) (mmol/L) represents the long-term average (i.e. daily) glucose concentration in the co-culture medium as given by:

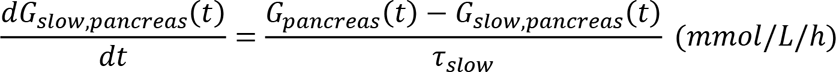

Insulin content in the liver compartment decreases over time due to insulin clearance by the liver spheroids (*CL*_*I,spheroids*_ (1/h)):

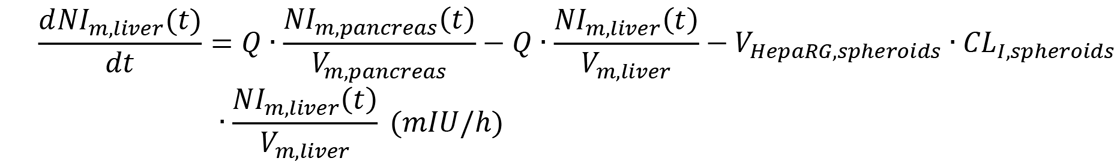

The concentrations of glucose and insulin in each compartment were calculated by dividing the insulin and glucose content, respectively, by the volume of co-culture medium in the compartment:

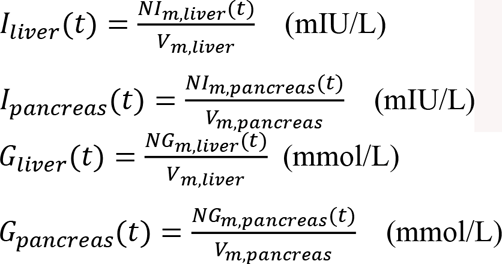

Glucose and insulin samples in the co-culture studies were obtained by pooling samples from both the liver and the pancreas compartment. Therefore, the resulting glucose and insulin measurements (*G*(*t*) and *I*(*t*), respectively), were computed as:

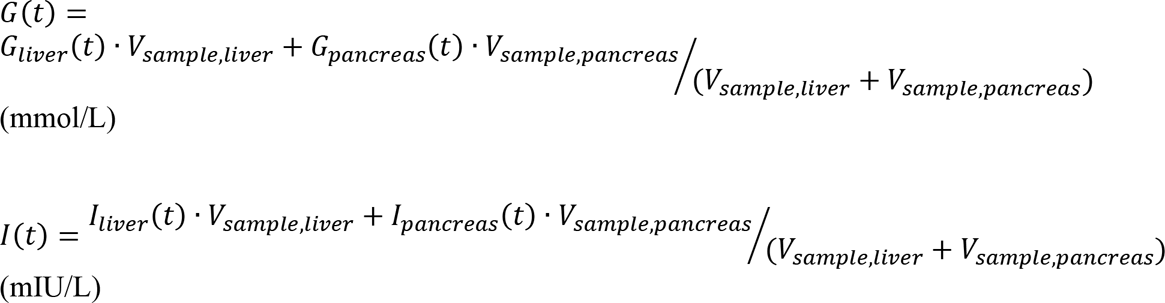

where *V*_*sample,liver*_ and *V*_*sample,pancreas*_ are the volumes of co-culture media collected from the liver and pancreas compartment, respectively, in each sample (15 µl).

The initial conditions for the model states are listed below:

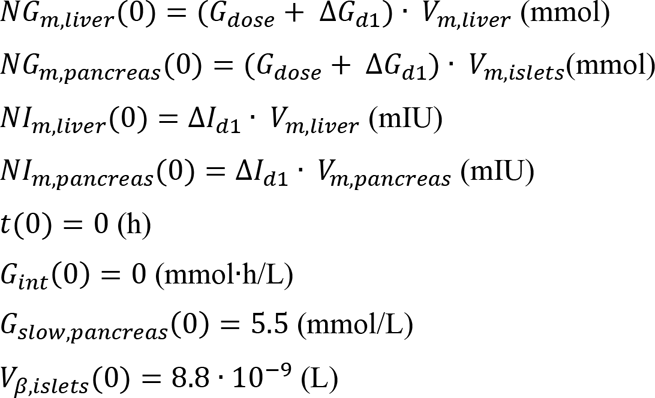

where Δ*G*_*d*1_(mmol/L), Δ*I*_*d*1_(*mIU*/*L*) are offset parameters that account for experimental errors related to the medium exchange performed on day 1. The experimental errors in the glucose concentration at 0 h can be due to varying co-culture medium volumes in the culture compartments, varying glucose concentration in the co-culture medium, or glucose assay-dependent variations. Values of insulin concentration different from zero at t=0 h could be due to co-culture medium remaining in the chip (both in the culture compartments and the microfluidic channel) during the medium exchange corresponding to the first GTT. Similarly, the model parameters (Δ*G*_*d*13_, Δ*I*_*d*13_) account for errors in glucose and insulin concentrations, respectively, during the medium exchanges performed on day 13.

#### Hypothesis testing to unravel the origin of insulin resistance in the pancreas-liver MPS

We tested two hypotheses that could explain the glucose and insulin responses observed in the MPS (**Fig. 1E**). The first hypothesis (H1) assumes that insulin resistance is caused by hyperglycemia alone, while the second hypothesis (H2) assumes that insulin resistance is caused by hyperglycemia and an additional diabetogenic factor. The model described in Casas et al.^11^ implements hypothesis H1. Therefore, we created a second computational model implementing hypothesis H2, by including an equation to model the effect of an additional diabetogenic factor on insulin sensitivity. This effect was modelled as a sigmoidal function of time, with maximal fractional reduction *I*_*max,additional*_, and with half of the maximal fractional reduction occurring at *EC*50_*additional*_ (mmol·h/L):

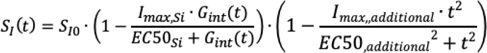

Each computational model was calibrated against the experimental data of glucose and insulin from the pancreas-liver co-culture. To perform this calibration, the model parameters were estimated using nonlinear optimization, by finding parameter values that provided an acceptable agreement with the experimental data according to the following cost function:

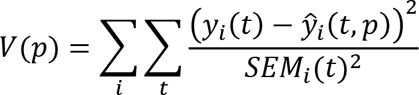

where *i* is summed over the number of experimental time-series for the given experiment *y*_*i*_(*t*) and 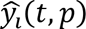 represents the model simulations and *p* the model parameters. SEM denotes the standard error of the mean and *t* the measured time points in each time-series. To handle uncertainty in the estimation, we used a simulated annealing approach^55^ to find the set of acceptable parameters that provided an acceptable agreement with the experimental data according to a statistical χ^2^ test^53, 56^ with a significance level of 0.05.

We found a good visual agreement with the experimental data for both models corresponding to H1 and H2 (**Fig. 2 C and E**). This visual agreement was statistically supported by the fact that both models passed a χ^2^ test at a significance level α = 0.05, with a value of the cost for the optimal parameter set *p*_*opt*_ lower than the χ^2^-threshold (*V*(*p*_*opt,H*1_) = 21.62 < 37.65, *V*(*p*_*opt,H*2_) = 28.32 < 37.65).

To be able to discriminate between H1 and H2, we performed predictions of glucose and insulin responses for different doses of added insulin to the co-culture medium, and selected an insulin dose that would provide detectable differences between the glucose responses for these hypotheses (i.e., differences larger than the average SEM across samples in the experimental data). The model predictions were made for the entire set of acceptable parameters. To visualize these predictions, we simulated model responses for the maximal and minimal values of each parameter within the set of acceptable parameters. We then calculated the boundaries of the prediction by computing the maximal and minimal value of the prediction for each time point and visualized the area between these boundaries (**Fig. 2C and D**). We performed the corresponding experiments for the calculated insulin dose (23 nM) and computed the model prediction. No acceptable agreement with the experimental data was found for H1, and this hypothesis was therefore rejected (**Fig. 2D**). H2, on the other hand, showed good visual agreement with the experimental data (**Fig. 2E**), which was also confirmed with a χ^2^ test at significance level α = 0.05 (*V*(*p*_*opt,H*2_) = 3.45 < 12.59).

#### Simulating the effect HCT concentration in the pancreas-liver MPS

In the computational model, the effect of high HCT on the pancreas-liver MPS was modelled as a decrease in both the insulin sensitivity of the liver spheroids *S*_*I*_(*t*) and the insulin secretion capacity of the *β* cells σ(*t*) over time, as follows:

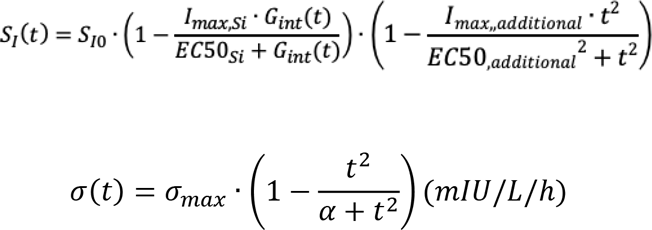

where the term 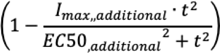 represents the effect of high HCT*S*_*I*_(*t*).

To model the effect of low HCT levels on the pancreas-liver MPS, we omitted the terms corresponding to these decreases in *S*_*I*_(*t*) and σ(*t*), leading to the following equations:

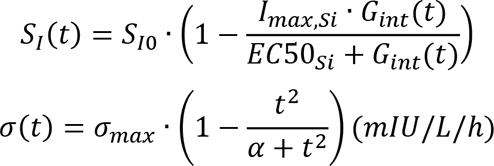

To predict the glucose and insulin responses in the pancreas-liver MPS under low HCT concentrations, we first calibrated the computational model using experimental data under high HCT levels from two GTT experiments, a GTT starting at day 1 (GTT day 1-3) and a GTT starting at day 13 (GTT day 13-15) (**Fig. 4 A**, high HCT). With the optimal parameter values obtained from this estimation as a start guess, we then optimized the parameters representing insulin sensitivity at the beginning of the co-culture *S*_*I*0_and the insulin secretion capacity of the beta cells using data under low HCT levels from a GTT starting at day 1 (**Fig 4A**, low HCT). We used this parameter set to predict the glucose and insulin responses under low HCT levels. In doing so, we omitted the decreases in *S*_*I*_(*t*) and σ(*t*) over time, as previously described.

#### Data pre-processing

Given the small number of replicate platforms in the MPS studies (two to six), we assume that the SEM values measured experimentally are an underestimation of the true uncertainty in the data. We considered SEM values below 5% of the corresponding mean to be unrealistic and corrected for possible measurement errors by setting these SEM values to the largest measured SEM value across all data points in the experimental dataset. Furthermore, we accounted for experimental errors in glucose and insulin measurements due to media-exchanges by including measured offsets in concentrations at the beginning of GTTs (t=0 within a given GTT) as an additional contribution to the total SEM for all data points corresponding to the given GTT. The resulting SEM values are given as error bars in all figures.

#### Software

Computations were carried out in MATLAB R2022b (The Mathworks Inc., Natick, Massachusetts, USA) using IQM tools (IntiQuan GmbH, Basel, Switzerland) and the MATLAB Global Optimization toolbox, as well as in Python (v 3.9.13). Figures 1A-C, 1E, 2A-C, and Figure S1 were prepared using BioRender (https://biorender.io/).

### Gene expression analysis

#### RNA isolation and quantitative real-time PCR

After the co-culture, liver spheroids in the culture compartments were washed three times with PBS and the spheroids were removed using a sterile blunt end needle (9180117, B.Braun) for RNA isolation. Spheroids were transferred into PCR-clean 1.5 mL microtubes with 100 µL of lysis buffer (LB1 from Macherey-Nagel or 700 µL of Buffer RLT (79216, Qiagen). Lysates were snap-frozen and stored at −80 °C. RNA was isolated using the NucleoSpin® RNA Plus XS kit (740990.50, Macherey-Nagel) or RNeasy Mini Kit (74104, Qiagen). cDNA was synthesized using TaqMan® Reverse Transcription Kit (Thermo Fisher Scientific). Real-time PCR was performed using the SensiFAST SYBR Lo-ROX Kit (BIO-94020, Bioline). Primers are shown in Supplementary Table 1. Relative gene expression was determined using the comparative CT (ΔΔCt) method with TBP as endogenous control gene.

#### RNA sequencing

The quantity and quality of RNA samples was assessed using the standard sensitivity RNA fragment analysis kit on Fragment Analyzer (Agilent Technologies). All samples had an RNA integrity number >8 and were deemed of sufficient quantity and quality for RNA-seq analysis. Samples were diluted to a final quantity of 150 ng/sample of total RNA. The KAPA mRNA HyperPrep kit (Roche) was used for reverse transcription, generation of double stranded cDNA and subsequent library preparation and indexing to facilitate multiplexing (Illumina TruSeq). All libraries were quantified with the Fragment Analyzer using the standard sensitivity NGS kit (Agilent Technologies) and pooled in equimolar concentrations and quantified with a Qubit Fluorometer (Thermo Fisher Scientific) with the DNA HS kit (Thermo Fisher Scientific). The library pool was further diluted to 2.2 pM and sequenced at >20M paired end reads/sample using the High Output regent kit to 150 cycles on an Illumina NextSeq500. RNASeq data was analysed using bcbio (version 1.1.0) and differential analysis was performed with DESeq2 (version 1.18.1).

### Proteomic analysis

#### Sample preparation for proteomic analysis

For proteomic analysis, the co-cultures were incubated in FBS-free co-culture medium for the last four days (d11-15). After finishing the culture, supernatants were collected from both pancreas and liver compartments and combined in a 1.5 mL microtube. Samples were first centrifuged at 300x*g* for 10 min at RT, to remove any remaining cells, and then supernatants were transferred into new tubes for centrifugation at 10,000x*g* for 10 min at 4 °C. The supernatants were stored at −80 °C until sample preparation for nano-scale liquid chromatographic tandem mass spectrometry (nLC-MS/MS) performed on two MPS media experiments and MS instruments, Q Exactive™ HF Orbitrap or Fusion™ Lumos™ Tribrid™ (Thermo Fisher Scientific).

#### Sample preparation, peptide labelling and fractionation for Q Exactive™ HF analysis

Equal volumes of the cell culture supernatants from each condition was concentrated on nanosep 10k omega filters (Pall Corporation, Port Washington, NY, USA) prerinsed with 50 mM triethylammonium bicarbonate (TEAB, Sigma-Aldrich) and was washed twice in the filter with 500 µL 50 mM TEAB, by spinning at 14,000×g for 20 min at 4 °C. Proteins were reduced on the filters using 100 µL 10mM TCEP (77720, Bond-Breaker™ TCEP solution, Thermo Scientific) in 50mM TEAB at 55 °C for 45 min followed by a 10 min spin at 14,000×g, 20°C. Free cysteine residues were modified using 100 µL freshly prepared 15 mM iodoacetamide (IAA, Sigma-Aldrich) in 50 mM TEAB and incubated for 20 min at room temperature in the dark. The IAA solution was removed by washing with 10% acetonitrile (ACN) in 50 mM TEAB followed by centrifugation and filters transferred to new LoBind Eppendorf tubes. Tryptic digestion was performed by adding 1.6 µg of trypsin (V5111, Promega, sequencing grade modified trypsin) in 40 µl 10% ACN in 50 mM TEAB and incubated at 37 °C under humid conditions. Next day digested peptides were collected after spinning and then rinsing the filters with 60 µL 10% acetonitrile in 50 mM TEAB followed by a final centrifugation at 14,000×g, which collected all tryptic peptides in the LoBind tube.

An equal amount (54 µg, determined by Pierce Quantitative Fluorometric peptide assay, 23275, Thermo Scientific) of peptides from each sample was subjected to isobaric labelling using Tandem Mass Tag (TMT-10plex) reagents (90110, Lot RG234662, Thermo Fischer Scientific) according to the manufacturer’s instructions. The labelled samples were combined into one pooled sample, concentrated using vacuum centrifugation and separated into eight fractions using Pierce™ High pH Reversed-Phase Peptide Fractionation Kit (84868, Thermo Scientific) according to the manufacturer’s instructions for TMT-labelled peptides. After vacuum centrifugation of peptide fraction to dryness, the peptides were resuspended in 0.2% Formic Acid (FA) in 3% ACN.

#### nLC-MS/MS with Q Exactive HF

The TMT-labelled peptide samples were analysed with an Easy-nLC1200 liquid chromatography system combined with Q Exactive HF mass spectrometer (Thermo Scientific) using a 136 min gradient. The separation was performed using an Acclaim PepMap precolumn (75 µM ID by 20 mm) connected to a 75 µM by 150 mm analytical Easy Spray PepMap RSLC C18 column (2µm particles, 100 Å pore size; Thermo Scientific) using a gradient from 5% solvent B to 15% solvent B over 47 min, then up to 25% B the next 58 min and up to 50% B in 20 min followed by an increase to 98% solvent B for 1 min, and 98% solvent B for 9 min at a flow of 280 nL/min. Solvent A was 0.1% formic acid and solvent B was 80% acetonitrile, 0.1% formic acid. MS scans were performed at 120 000 resolution, m/z range 350-1400. MS/MS analysis was performed in a data-dependent experiment, with top 15 of the most intense doubly or multiply positive charged precursor ions selected. Precursor ions were isolated in the quadrupole with a 1.2 m/z isolation window and 0.2 m/z offset, with dynamic exclusion set to a duration of 30 seconds. Isolated precursor ions were subjected to collision induced dissociation (CID) at 32 collision energy (arbitrary unit) with a maximum injection time of 100 ms. Produced MS2 fragment ions were detected at 60 000 resolutions, with a fixed first mass of 120 m/z and a scan range of 200-2000 m/z.

#### Proteomic Data Analysis of Q Exactive™ HF data

The data files were merged for identification and relative quantification using Proteome Discoverer version 2.1.1.21 (Thermo Fisher Scientific). Swiss-Prot Human database was used for the database search, using the Mascot search engine v. 2.5.1 (Matrix Science, London, UK) with MS peptide tolerance of 6 ppm and fragment ion tolerance of 0.02 Da. Tryptic peptides were accepted with 1 missed cleavage and methionine oxidation was set as a variable modification. Carbamidomethyl on cysteines and TMT on peptide N-termini and on lysine side chains were set as fixed modifications. Percolator was used for PSM validation with the strict FDR threshold of 1%. Quantification was performed in Proteome Discoverer 2.1.1.21. The TMT reporter ions were identified with 20 ppm mass tolerance in the MS2 spectra and the TMT reporter S/N values for each sample were normalized within Proteome Discoverer on the total peptide amount. Quantitative results were only based on unique peptide sequences with a co-isolation threshold of 50 and an average S/N threshold of 10 for the protein quantification.

#### Sample preparation, peptide labelling and fractionation for Fusion™ Lumos™ Tribrid™ analysis

Each sample was mixed with sodium dodecyl sulphate (SDS), triethylammonium bicarbonate (TEAB) and DL-dithiothreitol (DTT) to concentrations of 0.5% SDS, 50 mM TEAB, 100 mM DTT and incubated at 95 °C for 5 min for denaturation and reduction. The reduced samples were processed using the modified filter-aided sample preparation (FASP) method^57^. In short, the reduced samples were diluted to 1:4 by 8 M urea solution, transferred onto Nanosep 10k Omega filters (Pall Corporation, Port Washington, NY, USA) and washed repeatedly with 8 M urea and once with digestion buffer (0.5% sodium deoxycholate (SDC) in 50 mM TEAB). Free cysteine residues were modified using 10 mM methyl methanethiosulfonate (MMTS) solution in digestion buffer for 20 min at RT and the filters were washed twice with 100 µL of digestion buffer. One µg Pierce trypsin protease (MS Grade, Thermo Fisher Scientific) in digestion buffer was added and the samples were incubated at 37 °C for 3 hours. An additional portion of trypsin was added and incubated overnight.

The peptides were collected by centrifugation and isobaric labelling was performed using Tandem Mass Tag (TMT-10plex) reagents (Thermo Fischer Scientific) according to the manufacturer’s instructions. The labelled samples were combined into one pooled sample, concentrated using vacuum centrifugation, and SDC was removed by acidification with 10% TFA and subsequent centrifugation. The labelled pooled sample was treated with Pierce peptide desalting spin columns (Thermo Fischer Scientific) according to the manufacturer’s instructions.

Each purified desalted sample was pre-fractionated into 40 primary fractions with basic reversed-phase chromatography (bRP-LC) using a Dionex Ultimate 3000 UPLC system (Thermo Fischer Scientific). Peptide separations were performed using a reversed-phase XBridge BEH C18 column (3.5 μm, 3.0×150 mm, Waters Corporation) and a linear gradient from 3% to 40% solvent B over 18 min followed by an increase to 100% solvent B over 5 min and 100% solvent B for 5 min at a flow of 400 µL/min. Solvent A was 10 mM ammonium formate buffer at pH 10.0 and solvent B was 90% acetonitrile, 10% 10 mM ammonium formate at pH 10.0. The fractions were concatenated into 20 fractions, dried and reconstituted in 3% acetonitrile, 0.2% formic acid.

#### nLC-MS/MS with Fusion™ Lumos™ Tribrid™

The fractions were analysed on an orbitrap Fusion™ Lumos™ Tribrid™ mass spectrometer interfaced with Easy-nLC1200 liquid chromatography system (Thermo Fisher Scientific). Peptides were trapped on an Acclaim Pepmap 100 C18 trap column (100 μm x 2 cm, particle size 5 μm, Thermo Fischer Scientific) and separated on an in-house packed analytical column (75 μm x 35 cm, particle size 3 μm, Reprosil-Pur C18, Dr. Maisch) using a gradient from 5% solvent B to 33% solvent B over 77 min followed by an increase to 100% solvent B for 3 min, and 100% solvent B for 10 min at a flow of 300 nL/min. Solvent A was 0.2% formic acid and solvent B was 80% acetonitrile, 0.2% formic acid. MS scans were performed at 120 000 resolution, m/z range 375-1375. MS/MS analysis was performed in a data-dependent experiment, with top speed cycle of 3 s for the most intense doubly or multiply charged precursor ions. Precursor ions were isolated in the quadrupole with a 0.7 m/z isolation window, with dynamic exclusion set to 10 ppm and duration of 45 seconds. Isolated precursor ions were subjected to collision induced dissociation (CID) at 35 collision energy (arbitrary unit) with a maximum injection time of 50 ms. Produced MS2 fragment ions were detected in the ion trap followed by multinotch (simultaneous) isolation of the top 10 most abundant fragment ions for further fragmentation (MS3) by higher-energy collision dissociation (HCD) at 65% and detection in the Orbitrap at 50 000 resolutions, m/z range 100-500.

#### Proteomic Data Analysis of Fusion™ Lumos™ Tribrid™ data

The data files were merged for identification and relative quantification using Proteome Discoverer version 2.4 (Thermo Fisher Scientific). Swiss-Prot Human database was used for the database search, using the Mascot search engine v. 2.5.1 (Matrix Science, London, UK) with MS peptide tolerance of 5 ppm and fragment ion tolerance of 0.2 Da. Tryptic peptides were accepted with 0 missed cleavage and methionine oxidation was set as a variable modification. Cysteine methylthiolation and TMT on peptide N-termini and on lysine side chains were set as fixed modifications. Percolator was used for PSM validation with the strict FDR threshold of 1%. Quantification was performed in Proteome Discoverer 2.4. The TMT reporter ions were identified with 3 mmu mass tolerance in the MS3 HCD spectra and the TMT reporter S/N values for each sample were normalized within Proteome Discoverer 2.4 on the total peptide amount. Only the quantitative results for the unique peptide sequences with the minimum SPS match % of 40 and the average S/N above 10 were included for the protein quantification.

Proteomic data was compared to RNASeq results by pairing log2 fold-changes at the gene level and plotted in Fig. 6B. Data was plotted with R version 4.0.2 with ggplot2 version (3.3.5).

### Lipid vesicle staining

Liver spheroids in the culture compartments were fixed with 4% methanol-free paraformaldehyde (PFA; 28908, Thermo Scientific) at 4 °C overnight. On the following day, the spheroids were washed three times with PBS and then stored in PBS at 4 °C until use. Fixed spheroids were stained with 2 µM Nile Red (72485, Sigma-Aldrich) and 16 µM Hoechst 33342 (Invitrogen) in PBS. Samples were first incubated at 37 °C for 2 hours, followed by an overnight incubation at RT. Next, the staining solution was removed, and the compartments were washed three times with PBS. Fluorescence imaging was performed using confocal laser scanning microscope (LSM880 Airyscan Zeiss,) and image processing and reconstruction were carried out using ZEN 3.2 software (Zeiss).

### Glycogen staining

Liver compartments were washed with 0.1% BSA in PBS and the liver spheroids were detached from the bottom of the culture compartment using a sharp needle. The spheroids were transferred into 1.5 ml microtubes using wide-bore pipette tips for fixation using 4% methanol-free PFA at 4°C overnight. The spheroids were then washed three times with PBS and stored at 4 °C until use. PAS staining to visualize the storage of glycogen was performed by Histocenter (Mölndal, Sweden). Briefly, after standard paraffin embedding and sectioning, the sections were sequentially treated with 0.5% periodic acid, water, Schiff reagent, water, Weigert’s iron haematoxylin solution, water, hydrochloric acid, water, and 95% ethanol. Imaging was carried out using an inverted microscope (Axiovert 40 CFL, Zeiss).

### Cell proliferation analysis by EdU incorporation

We developed a method to quantify cell proliferation in pancreatic islets by using EdU incorporation, automated HT imaging, optical slicing, and automated image analysis (**Fig. S8A**). To test robustness of the method, islets were cultured in Akura™ 96 Spheroid Microplate for 4 days, either in Human Islet Maintenance Medium (untreated control) or in the presence of 10 µM of the MST1 kinase inhibitor 4-(5-amino-6-(1-oxo-1,2,3,4-tetrahydroisoquinolin-6-yl)pyrazin-2-yl)-N-cyclopropyl-N-methylbenzenesulfonamide^58^ (CAS 1396771-17-7) which was used as a positive control. To label proliferating cells, media were supplemented with 10 µM EdU. Donor for the robustness analysis study was a male, 45 years with BMI of 29.8 and HbA1c of 5.10%.

Fixation, permeabilization, and EdU staining were performed in Akura™ 96 Spheroid Microplates. The islets were fixed with 4% PFA at RT for 2 hours, washed twice with 0.1% BSA in PBS, and permeabilized with 1x BD Perm/Wash buffer (554723, BD Biosciences) for 1 hour at RT. Next, the islets were stained with Click-iT EdU reaction cocktail (C10638, Click-iT® Plus EdU Alexa Fluor® 555 Imaging Kit, Molecular Probes), for 2 hours at RT in dark. After removal of the reaction cocktail, islets were washed once with 1x BD Perm/Wash buffer and transferred into Akura™ 384 Spheroid Microplate (CS-09-003-02, InSphero). Finally, a sorbitol-based clearing reagent Sca*l*e S4(0)^59^ (40 (w/v)% D-(-)-Sorbitol (S3889, Sigma-Aldrich), 10(w/v)% Glycerol (G9012, Sigma-Aldrich), 4 M Urea (U0631, Sigma-Aldrich), 15-25(v/v)% DMSO) containing 3.0-3.9 μM SiR-DNA^60^ (Spirochrome) for nuclear staining was added, and the plate was incubated overnight at RT. The plate was then centrifuged at 700x*g* for 1 min to remove bubbles and collect islets in the middle of wells, and stored at 4 °C until imaging.

Images were acquired on a CellVoyager 7000 high-throughput spinning disc confocal microscope (Yokogawa). All microwells were first screened using a 10X 0.16NA objective at 2×2 binning (**Fig. S8**B). A MATLAB-based Search First script (Wako Software Suite; Wako Automation) was used for automated detection of islet position in each micro-well. Then, high-resolution z-stacks of 200 µm from well bottom were acquired for each islet at its exact position, using a 40X 0.75NA objective at 2×2 binning in 2 fluorescence channels – EdU-positive nuclei (Click-iT EdU Alexa Flour 555; 561 nm laser) and nuclei (SiR-DNA far-red DNA stain; 640 nm laser) (**Fig. S8C**). Using optical clearing in combination with 561 nm and 640 nm laser allowed for penetration of laser light and acquisition of fluorescent signal from throughout the islets, which usually have a diameter of 100 - 150 µm. Analysis of total number of nuclei and EdU-positive nuclei was performed using Columbus™ Image Data Storage and Analysis system (ver. 2.8.1, Perkin Elmer).

We observed that the percentage of EdU positive cells is largely independent of optical sampling distance in the range of 0.4-20 µM (**Fig. S8D**). Islets treated with the MST1 kinase inhibitor showed a significantly higher number of EdU positive cells as compared to the untreated control islets (**Fig. S8E**) demonstrating that the developed method can reliably separate different study groups.

In the pancreas-liver co-cultures, 10 µM EdU was added into co-culture medium for the last 5 days to label proliferating cells. After finishing a co-culture, islets were first transferred from chips into individual wells of an Akura™ 96 Spheroid Microplate followed by fixation, permeabilization, staining, and imaging as described above.

### Statistical analysis

GraphPad Prism software (Version 8) was used to plot the data and perform comparative analysis between the means of different conditions. For comparing two unpaired means of normal distributed data with homogenous variance, a two-tailed Student’s t-test or a multiple t-test using the Holm-Sidak method (in case of several independent comparisons e.g., for comparing gene expression of multiple genes between two conditions) was performed. Normality was tested using the Shapiro-Wilk normality test and equality of variance was tested using the F-test. A p value < 0.05 was considered statistically significant.

For comparing three or more means of normal distributed data with homogenous variance, a one-way ANOVA was performed. Normality was tested using the Shapiro-Wilk normality test and equality of variance was tested using the Brown-Forsythe test. Bonferroni’s multiple comparison post-hoc test was used to compare the means of several conditions to a control mean and Sidak’s multiple comparison post-hoc test was used to compare the mean of selected pairs of conditions. A p value < 0.05 was considered statistically significant.

Fold changes of gene expression in Fig. 3 and 5 and GSIS data in Fig. 3E were log-transformed for normality. The area under the glucose and insulin GTT curves was calculated with Prism using the trapezoidal method.

## ACKNOWLEDGEMENTS

We thank S. Prill, A. Vildhede, B. Nugraha, N. Nguyen, K. Schimek, T. P. Tao, I. Durieux, S. Aljburi, and I. Rütschle for technical assistance. Part of the proteomic analysis was performed at the Proteomics Core Facility at Sahlgrenska Academy, University of Gothenburg. PAS staining was performed at Histocenter AB, Mölndal, Sweden

## DATA AVAILABILITY

The code for data analysis, visualization, and mathematical modelling is publicly available on <u>GitHub</u>.

