## Supplementary Information for "Diseased human pancreas and liver microphysiological system for preclinical diabetes research"

Sophie Rigal *et al.*

**This PDF file includes:**

Figs. S1 to S10
Table S1

**Fig. S1.**

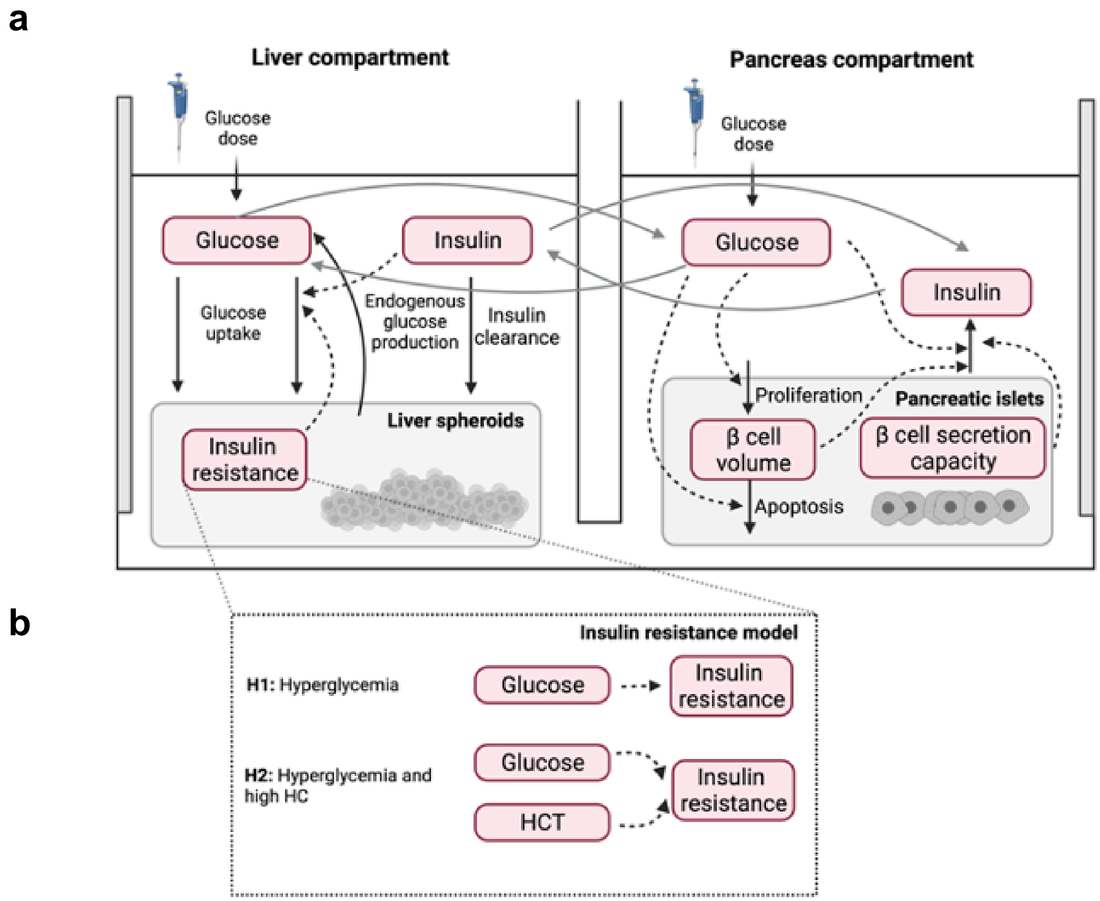


**Overview of the mathematical model to study the development of insulin resistance in the pancreas-liver co-culture. (A)** Previously reported mathematical model describing glucose metabolism in the pancreas-liver co-culture[11]. The model describes the behavior of key physiological variables involved in glucose homeostasis (red text boxes) and includes both liver (left) and pancreas (right) compartments, which represent the corresponding co-culture compartments in the MPS. The dashed arrows represent the interactions between the physiological variables described in the model, and the solid arrows represent metabolic fluxes in the co-culture. **(B)** Implementation of the hypotheses to study the development of insulin resistance. In the first hypothesis (H1), the insulin resistance variable is determined by the glucose content in the co-culture medium. In the second hypothesis (H2), insulin resistance is dependent on both glucose content and the concentration of hydrocortisone (HCT).

**Fig. S2.**

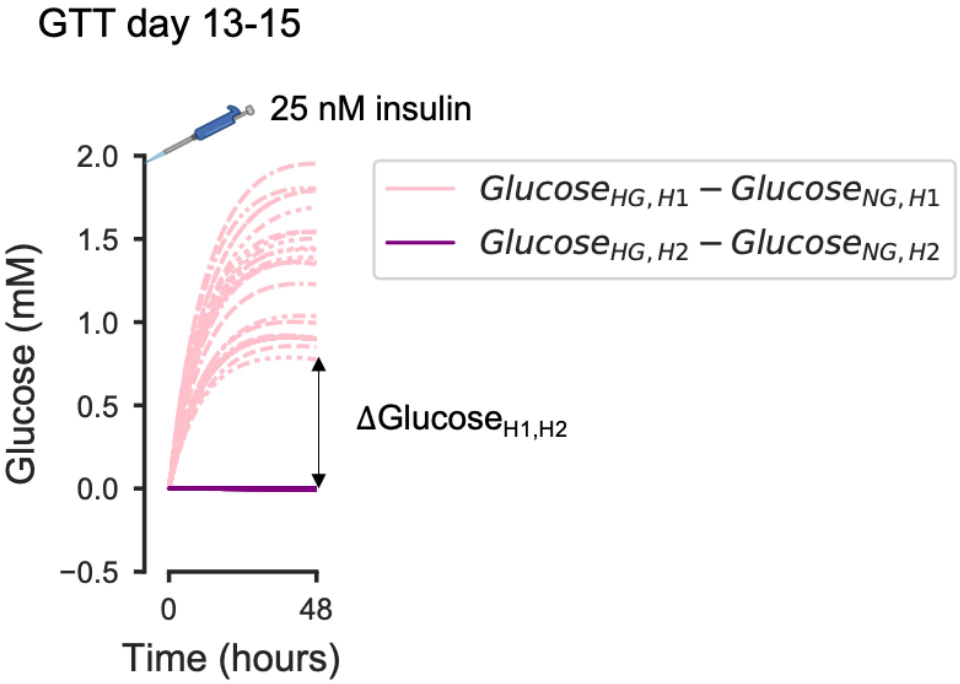


***In silico*-based calculation of the insulin dose to spike into the co-culture for differentiating between hypotheses.** We used the mathematical model calibrated using data from GTTs on days 1-3 and 7-9 to predict the effect of an insulin dose on glucose regulation for the studied hypothesis H1 and H2 (Fig. 2b). The lines represent model-predicted differences in glucose concentration between hyper- and normoglycemic co-cultures during the GTT on days 13-15 for both hypotheses H1 (pink lines) and H2 (purple lines) when a certain insulin dose is spiked into the co-culture medium. We simulated these glucose concentrations for a range of insulin doses using the mathematical model and selected the dose that yielded a concentration difference between the hypotheses ($G_{H1,H2}$) larger than the average SEM across all glucose experimental measurements on days 1-3 and days 7-9 (0.48 mM). The simulations shown in the figure correspond to the selected insulin dose (25 nM), where each line represents a model prediction with an acceptable parameter set.

**Fig. S3.**

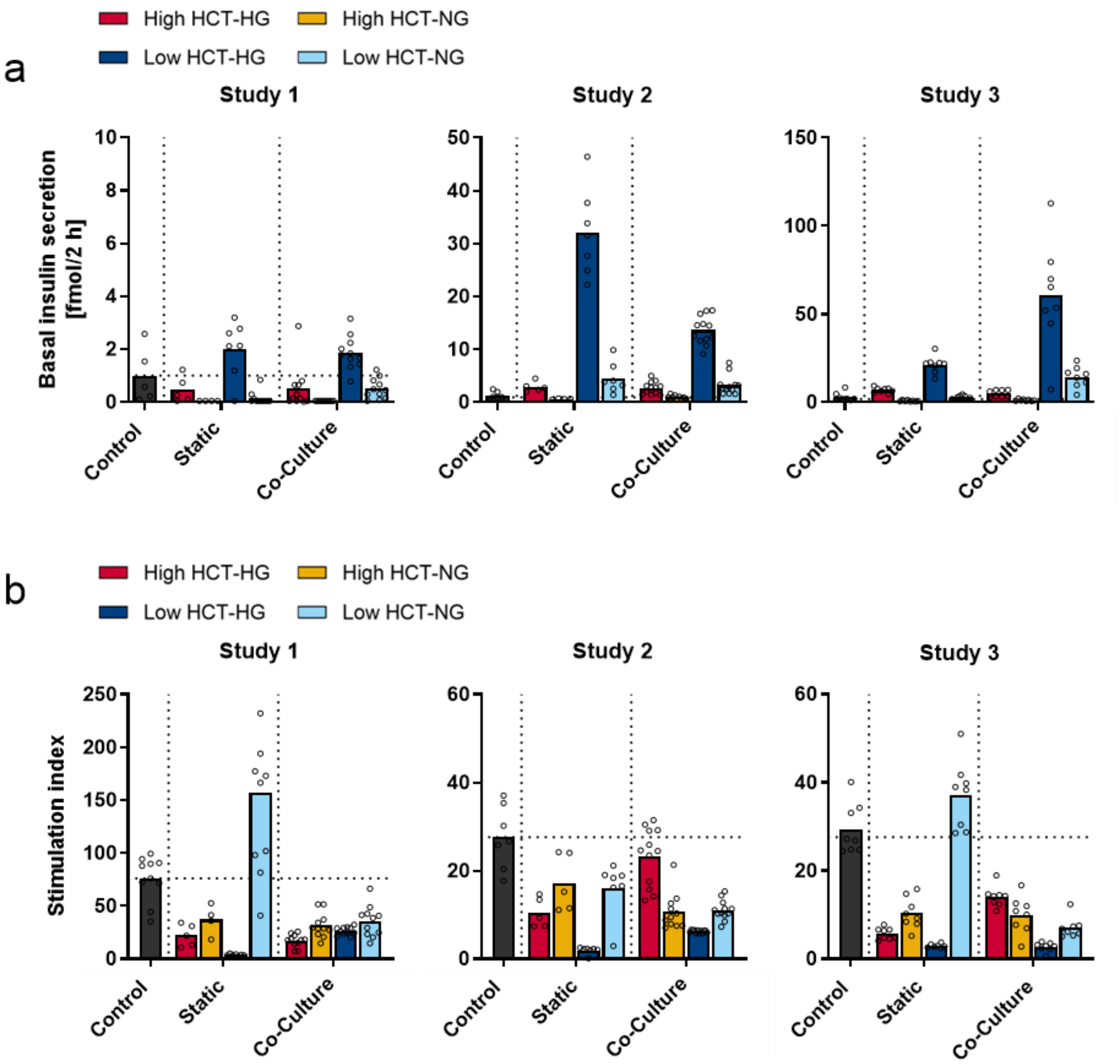


**Effect of glucose and HCT on islets.** (**A**) GSIS response after low-glucose stimulation (2.8 mM) in islets cultured for 15 days in static or in chip co-culture. Data shown as a fold change to static islets cultured in the supplier's maintenance medium that served as a control. (**B**) GSIS stimulation index showing the high glucose buffer (16.8 mM) increased insulin secretion relative to low-glucose buffer (2.8 mM). Symbols represent individual islets, an individual donor was used for each study (**A, B**).

**Fig. S4.**

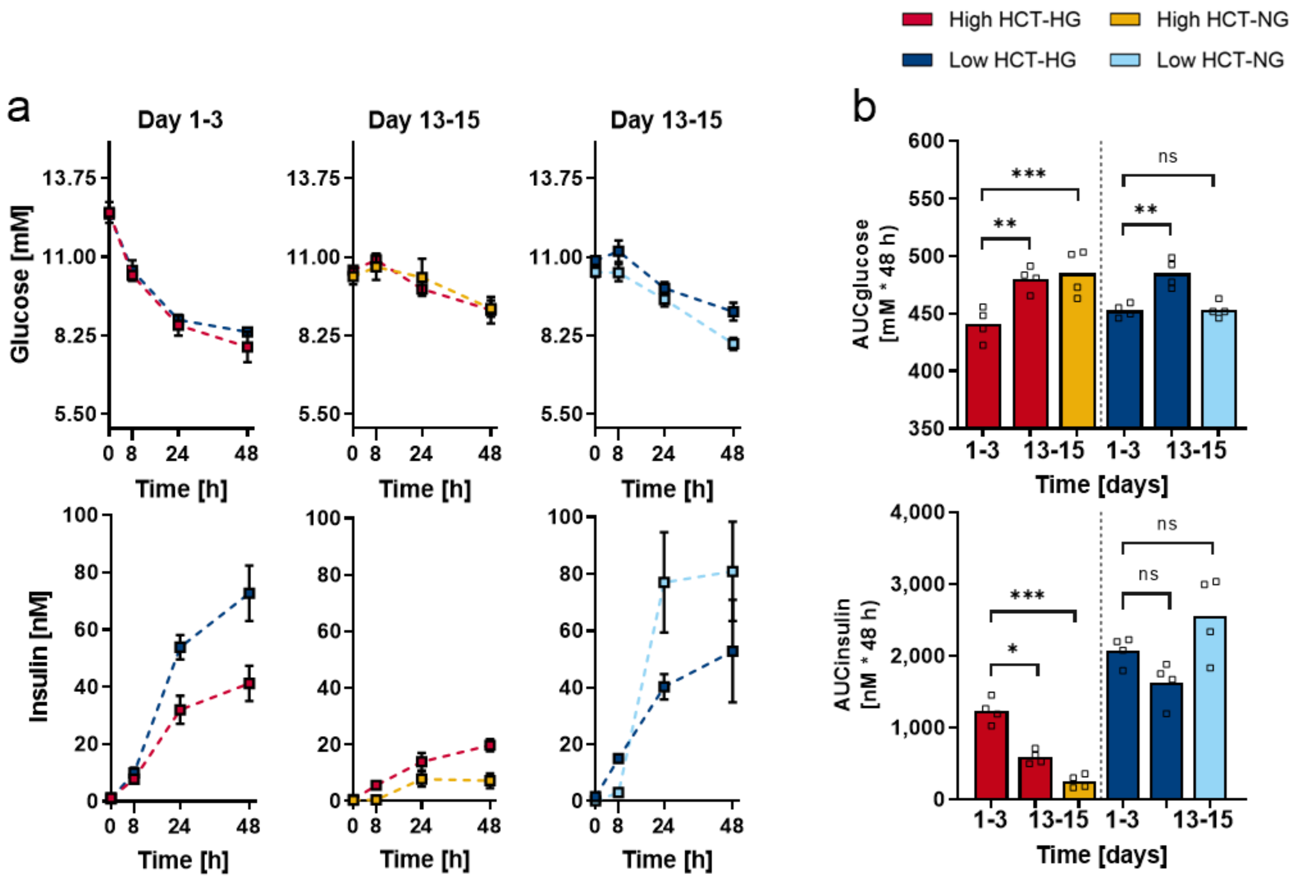


**Effect of glucose and HCT on glucose and insulin responses.** (**A**) Glucose and insulin concentration during glucose tolerance test. (**B**) Area under the curve (AUC) for glucose (left) and insulin (right). Symbols represent individual pancreas-liver co-cultures from one study. Differences between selected pairs of conditions (day 13-15 compared to day 1-3) were evaluated by one-way ANOVA using Sidak's multiple comparisons post-hoc test, ns = not significant, *p < 0.05, **p < 0.01, ***p < 0.001.

**Fig. S5.**

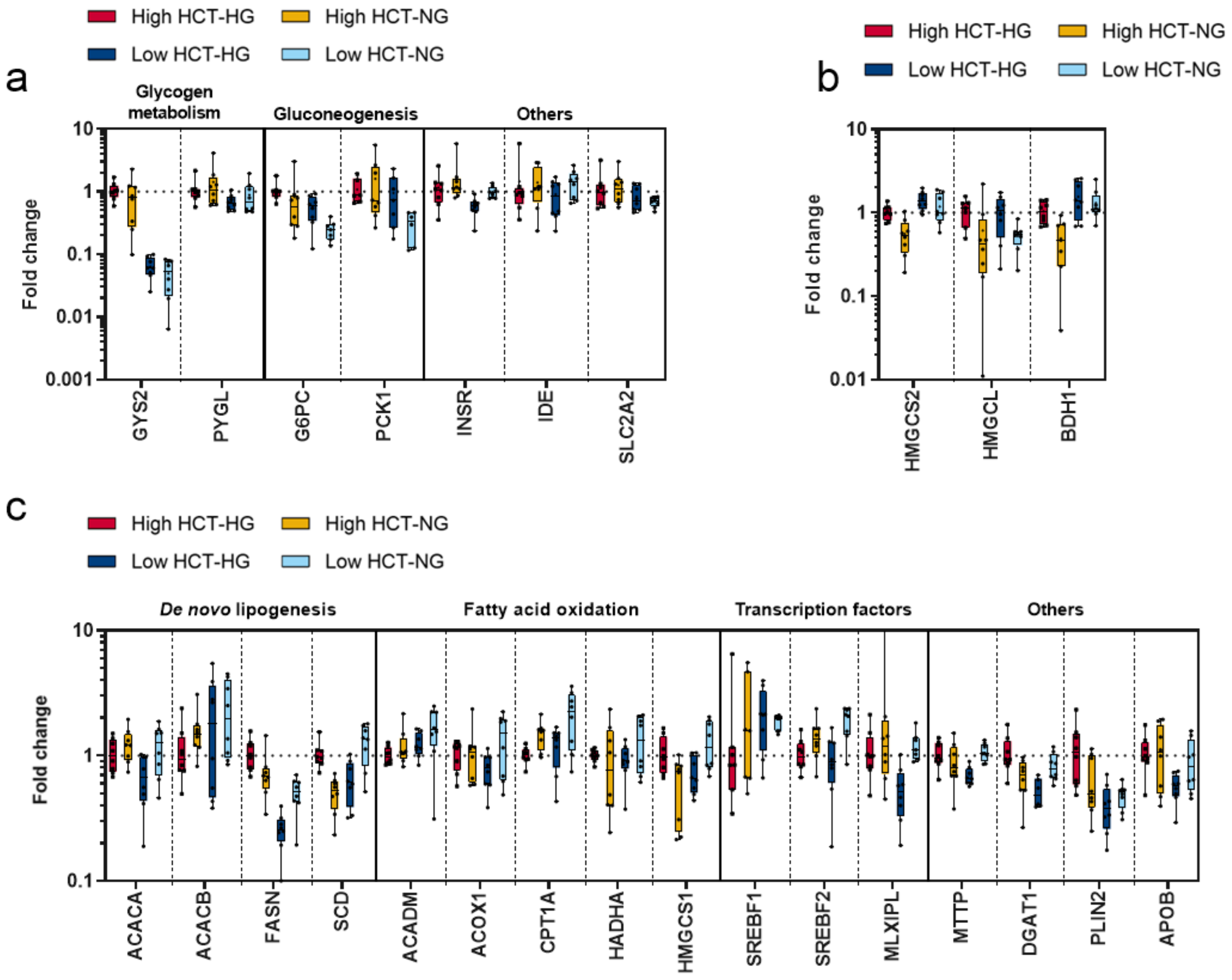


**Effect of glucose and HCT on gene expression in HepaRG/HHSteC liver spheroids. (A, B, C)** Gene expression of enzymes involved in hepatic glucose metabolism (**A**), ketone body synthesis (**B**), and lipid metabolism (**C**). Data shown as fold change between diseased and healthy condition in a box-whisker plot with geometric mean and min-max values. Symbols represent replicates from two independent studies, total n = 8.

**Fig. S6.**

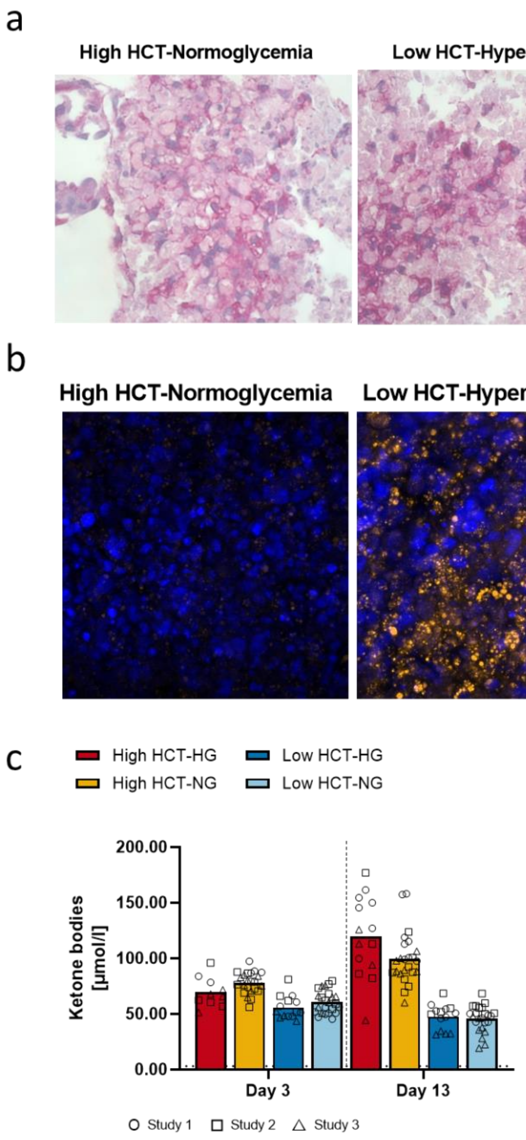


**Characterization of insulin resistance-associated phenotype of HepaRG/HHSteC liver spheroids. (A)** Glycogen storage visualized by periodic acid-Schiff (PAS) staining. (**B**) Intracellular lipid vesicles visualized by Nile Red staining (amber colour). Blue denotes DAPI-stained nuclei. (**C**) Ketone body synthesis represented by 3-hydroxybutyrate concentration in the co-culture supernatants. Symbols represent co-culture replicates from three independent studies.

**Fig. S7.**

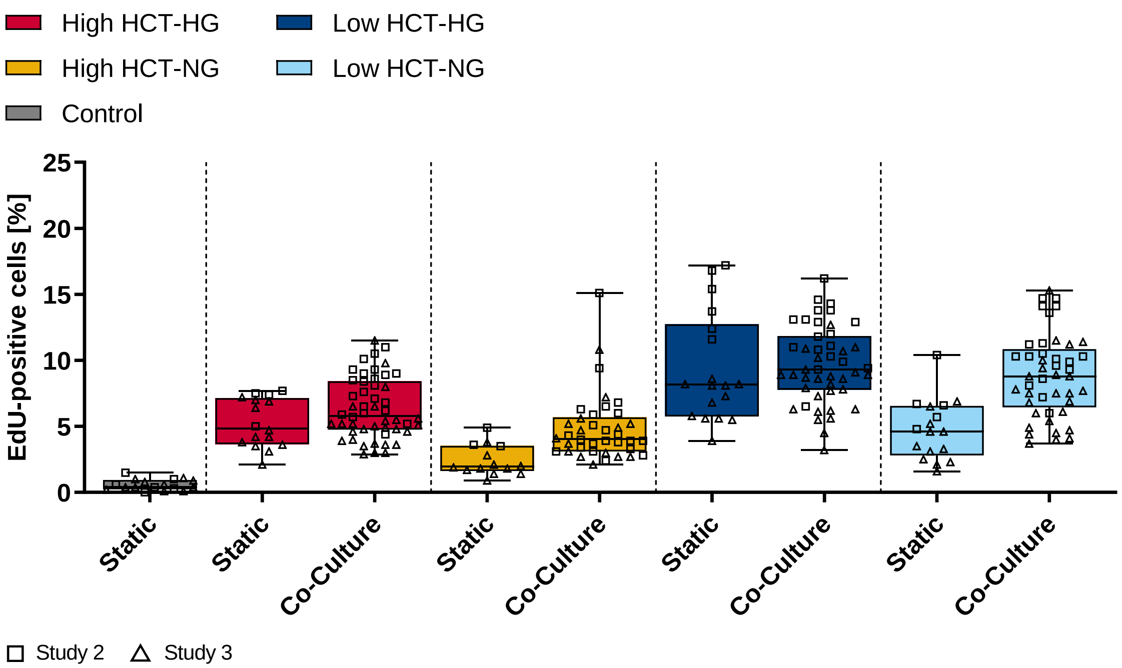


**Islet proliferation.** Proliferation of islets at the end of the culture as analysed by the percentage EdU-positive cells. Squares (study 2) and triangles (study 3) represent individual islets from two independent co-culture studies.

**Fig. S8.**

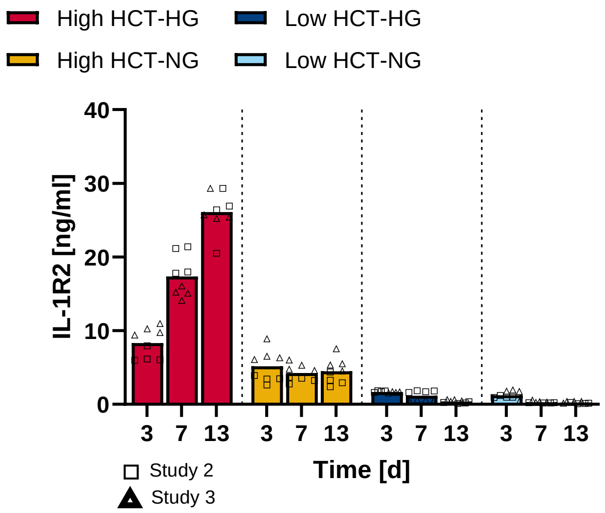


**Secretion of IL-1R2.** IL-1R2 concentration in the chip co-cultures over time. Squares (study 2) and triangles (study 3) represent individual islets from two independent co-culture studies, total n = 8.

**Fig. S9.**

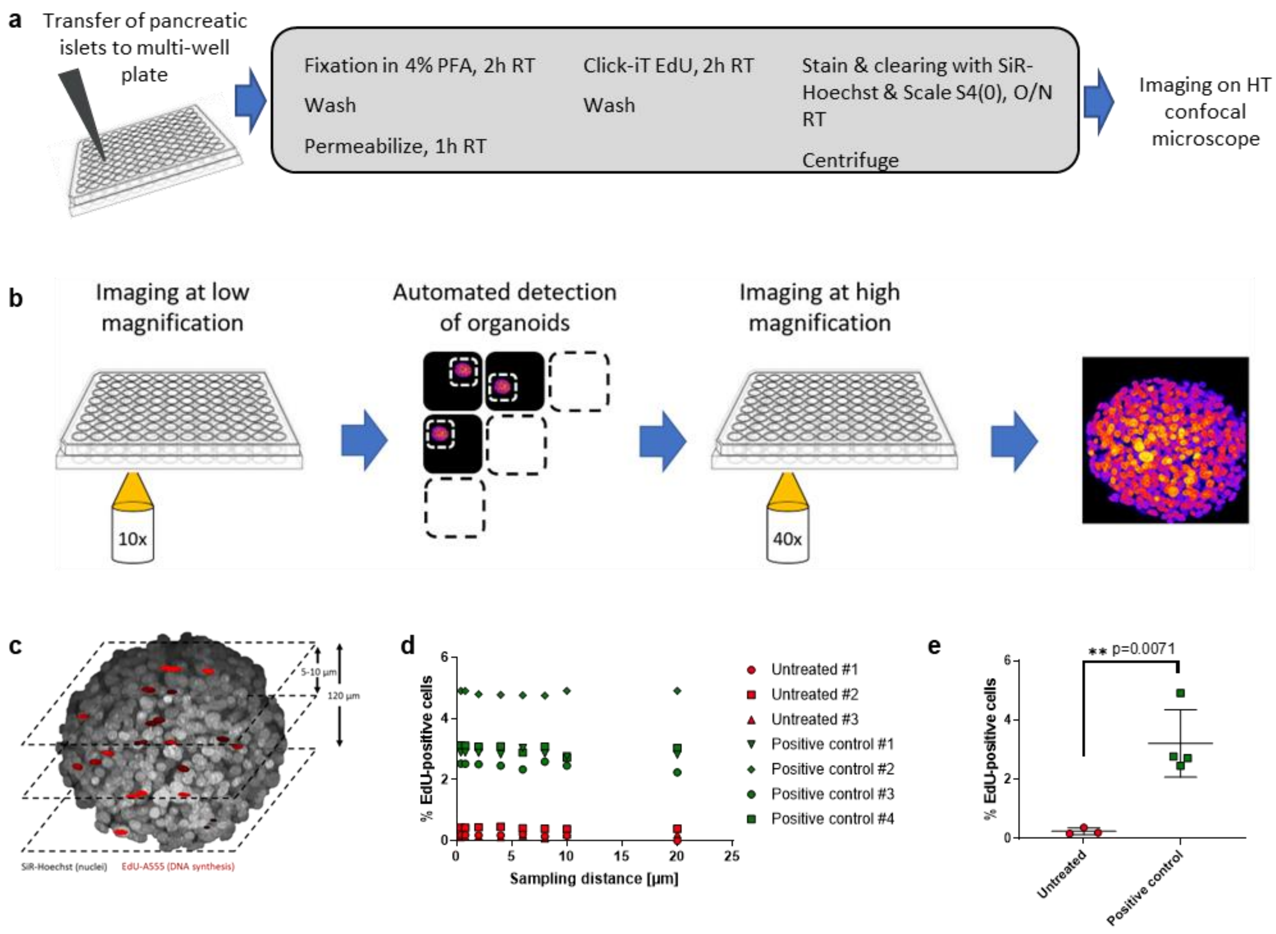


**Robust detection of cell proliferation in human pancreatic islets using EdU incorporation, automated high-throughput confocal microscopy, and optical slicing.** (**A**) Method description for pancreatic islet preparation, clearing, and staining for high-throughput imaging. (**B**) Schematic of automated detection of islets for high resolution imaging using Search First algorithm in multi-well plates. (**C**) Schematic of optical slicing of islets by confocal microscopy. (**D**) The percentage of EdU-positive cells is largely independent of the sampling distance (range 0.4–20 µm) as shown with untreated islets and islets treated with MST1 kinase inhibitor[58]. (**E**) The percentage of EdU-positive cells in untreated islets and in islets treated with MST1 kinase inhibitor. Data represents mean and SD. Significance determined by unpaired t-test.

**Fig. S10.**

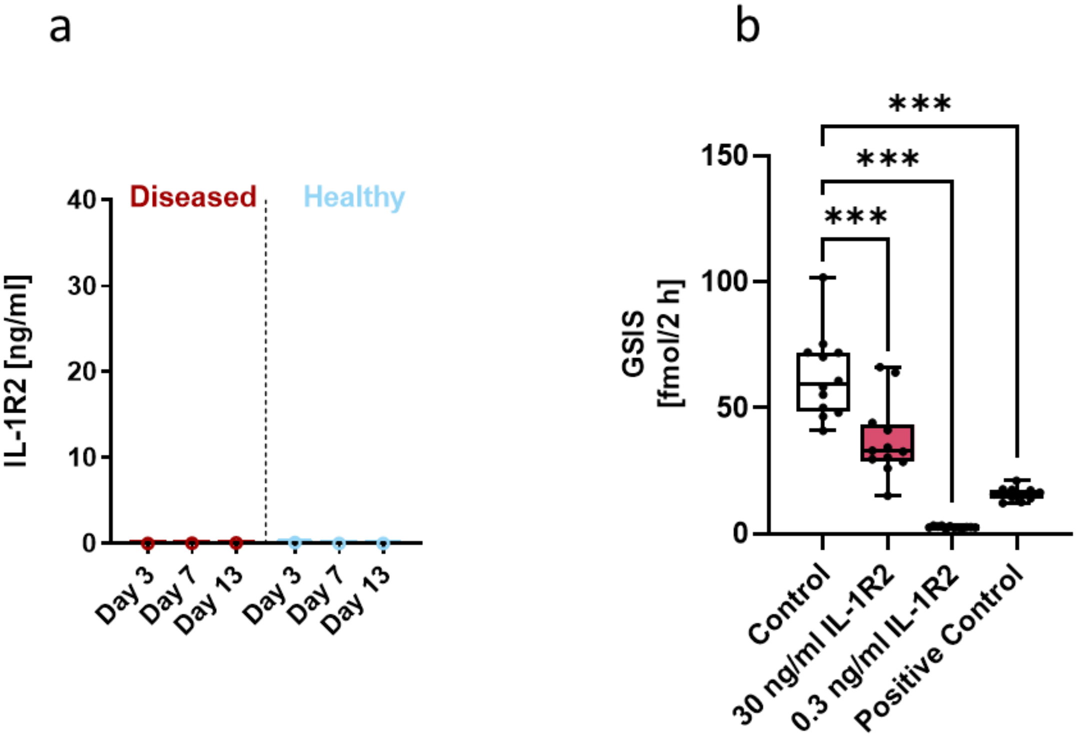


**Secretion and effect of IL-1R2. (A)** IL-1R2 concentration in static islet mono-cultures in the diseased (11 mM glucose, 50 µM HCT) and healthy (5.5 mM glucose, 10 nM HCT) conditions over time. Symbols represent individual islets. (**B**) IL-1R2 treatment decreases glucose stimulated insulin secretion (GSIS) both at low (0.3 ng/mL) and high (30 ng/mL) IL-1R2 dose in islets mono-cultured in static condition in Human Islet Maintenance Medium (InSphero). Data represents GSIS response after high-glucose stimulation (16.8 mM). Control, 0 ng/mL IL-1R2. Positive control, low hydrocortisone-normoglycemic (healthy) co-culture medium. Symbols represent individual islets. Differences to the control were evaluated by one-way ANOVA using Dunnett’s multiple comparisons post-hoc test, ***$p < 0.001$.

**Table S1.**
**Primers used in the qPCR.**

| Gene symbol | Name | Forward primer (5‘ - 3‘) | Reverse primer (5‘ - 3‘) |
|---|---|---|---|
| ACACA | Acetyl-CoA carboxylase alpha | AATAAGGATCTGGCGGAGTGG | GCTCGCTGAGTGGGTGATATG |
| ACACB | Acetyl-CoA carboxylase 2 | GCCGACTTCCATGACACACC | TGTTCCAGCCACTGCACAAC |
| ACADM | Acyl-Coenzyme A dehydrogenase | TGTTTTAATTGGTGACGGAGCTG | ACCAGAATCAACCTCCCAAGC |
| ACOX1 | Acyl-CoA oxidase 1 | GCAAGGAGGTAGCTTGGAACC | GATGCTCCCCTGAAGGAAATC |
| ACTA2 | Alpha-actin-2 | AGAGACCCTGTTCCAGCCATC | CGTGATCTCCTTCTGCATTCG |
| AHSG | Alpha 2-HS glycoprotein | CGCAAAATGTGATTCCAGTCC | CCGTTGTTCTGAGCGTTGAAG |
| ALB | Albumin | TCAGCTCTGGAAGTCGATGAAAC | AGTTGCTCTTTTGTTGCCTTGG |
| APOB | Apolipoprotein B | AGGCATCTCCACCTCAGCAG | GCTGCCTCTTCTTCCCAATTAAC |
| BDH1 | 3-hydroxybutyrate dehydrogenase 1 | AGCAGGTGGCAGAAGTGAACC | CTACCCCGAACTTGGTGATGC |
| BSEP/ABCB11 | Bile salt export pump/ATP binding cassette subfamily B member 11 | GCAGACACTGGCGTTTGTTG | ATGTTTGAGCGGAGGAACTGG |
| CPS1 | Carbamoyl-phosphate synthase 1 | CCCAGCCTCTCTTCCATCAG | GCGAGATTTCTGCACAGCTTC |
| CPT1A | Carnitine palmitoyltransferase 1A | CGTCACCTCTTCTGCCTTTACG | CTCCGCTGGACACGTACTCTG |
| CYP3A4 | Cytochrome P450 family 3 subfamily A member 4 | GGAAGTGGACCCAGAAACTGC | TTACGGTGCCATCCCTTGAC |
| DGAT1 | Diacylglycerol O-Acyltransferase 1 | GGCCTTACCTGGCTACACTGG | CATTGCCACTCCCATTCTTTG |
| FASN | Fatty acid synthase | ACGGACATGGAGCACAACAG | GGTACTTGGCCTTGGGTGTG |
| G6PC | Glucose-6-phosphatase catalytic subunit | GCTGAATGTCTGTCTGTCACGAA | CGAAGCTGAACAGGAAGAAGGT |
| GYS2 | Glycogen synthase 2 | GCCAGACACCTGACATTAAGCA | TGAGACCCTGAAGGAGAAGGTG |
| HADHA | Hydroxyacyl-CoA dehydrogenase trifunctional multienzyme complex subunit alpha | TGGTAGAAGCATTCGTGCAGAC | GGCATACGCTGTCAATTTTTCC |
| HMGCL | 3-hydroxy-3-methylglutaryl-CoA lyase | TCTACTCAATGGGCTGCTACGAG | TCCTGCCACAGAAGAGTCCAC |
| HMGCS1 | 3-hydroxy-3-methylglutaryl-CoA synthase 1 | TTGAGTCCAGCTCTTGGGATG | CACTGGGCATGGATCTTTTTG |
| HMGCS2 | 3-hydroxy-3-methylglutaryl-CoA synthase 2 | CCACCACTCTGCCCAAGAAC | ACACACTTTCGGGAGGCTAGG |
| HNF4 | Hepatocyte nuclear factor 4 alpha | ATACGCATCCTTGACGAGCTG | CTGGCGGTCGTTGATGTAGTC |
| IDE | Insulin degrading enzyme | TGGATTCTTGTCCTGTTGTTGG | TCAGAGTTTTGCAGCCATGAAG |
| INSR | Insulin receptor | CACGCCAAAGGACCACATGTC | CGAGAAACTGCATGGTCGCC |
| MLXIPL | MLX interacting protein like | AGCTGCGGGATGAGATTGAG | TCAAACAGAGGCCGGATGAG |
| MRP2/ABCC2 | Multidrug resistance-associated protein 2/ATP binding cassette subfamily C member 2 | GCATCCACAGACATCAGGTTCAC | CTGCGGCTCTCATTCAGTCTTTC |
| MTTP | Microsomal Triglyceride Transfer Protein | TGCAATGGAGTTTAGCTTGTGG | GTTTCCAGGCCAGCTTTCAC |
| PCK1 | Phosphoenolpyruvate carboxykinase 1 | CATGAGATCAGAGGCCACAGC | AATTTGCCTTCCTTGTCCTTCC |
| PLIN2 | Perilipin 2 | TTCACTCCCGTGCCTACCAG | CATCATATCCAATGCTCCTTTTCC |
| PYGL | Glycogen phosphorylase L | GATGTGGCTGCTTTGGACAAG | TCTTGACACTTGACATAGGCTTCG |
| SCD | Stearoyl-CoA desaturase | GCCACCGCTCTTACAAAGCTC | CGTCGGGAATTATGAGGATCAG |
| SLC2A2 | Solute carrier family 2 member 2/GLUT2 | TGGTTCATGGTGGCTGAGTTT | AGGGTAAAGGCCAGGAGCAC |
| SREBF1 | Sterol regulatory element binding transcription factor 1 | TCTGAGAGACCCCTGCCCAG | CTGCACGGCCTTGTCAATGG |
| SREBF2 | Sterol Regulatory Element Binding Transcription Factor 2 | GCTGTGCGCTCTCATTTTACC | GACGTTGAGGCTGCTCCATAG |
| TBP | TATA-box binding protein | CCTTGTGCTCACCCACCAAC | TCGTCTTCCTGAATCCCTTTAGAATAG |
